# The Conformational Plasticity of the Selectivity Filter Methionines Controls the In-Cell Cu(I) Uptake through the CTR1 transporter

**DOI:** 10.1101/2021.11.04.467269

**Authors:** Pavel Janoš, Jana Aupič, Sharon Ruthstein, Alessandra Magistrato

## Abstract

Copper is a trace element vital to many cellular functions. Yet its abnormal levels are toxic to cells, provoking a variety of severe diseases. The high affinity Copper Transporter 1 (CTR1), being the main in-cell copper (Cu(I)) entry route, tightly regulates its cellular uptake via a still elusive mechanism. Here, all-atoms simulations unlock the molecular terms of Cu(I) transport in eukaryotes disclosing that the two Methionine triads, forming the selectivity filter, play an unprecedented dual role both enabling selective Cu(I) transport and regulating its uptake-rate thanks to an intimate coupling between the conformational plasticity of their bulky side chains and the number of bound Cu(I) ions. Namely, the Met residues act as a gate reducing the Cu(I) import-rate when two ions simultaneously bind to CTR1. This may represent an elegant autoregulatory mechanism through which CTR1 protects the cells from excessively high, and hence toxic, in-cell Cu(I) levels. Overall, these outcomes resolve fundamental questions in CTR1 biology and open new windows of opportunity to tackle diseases associated with an imbalanced copper uptake.

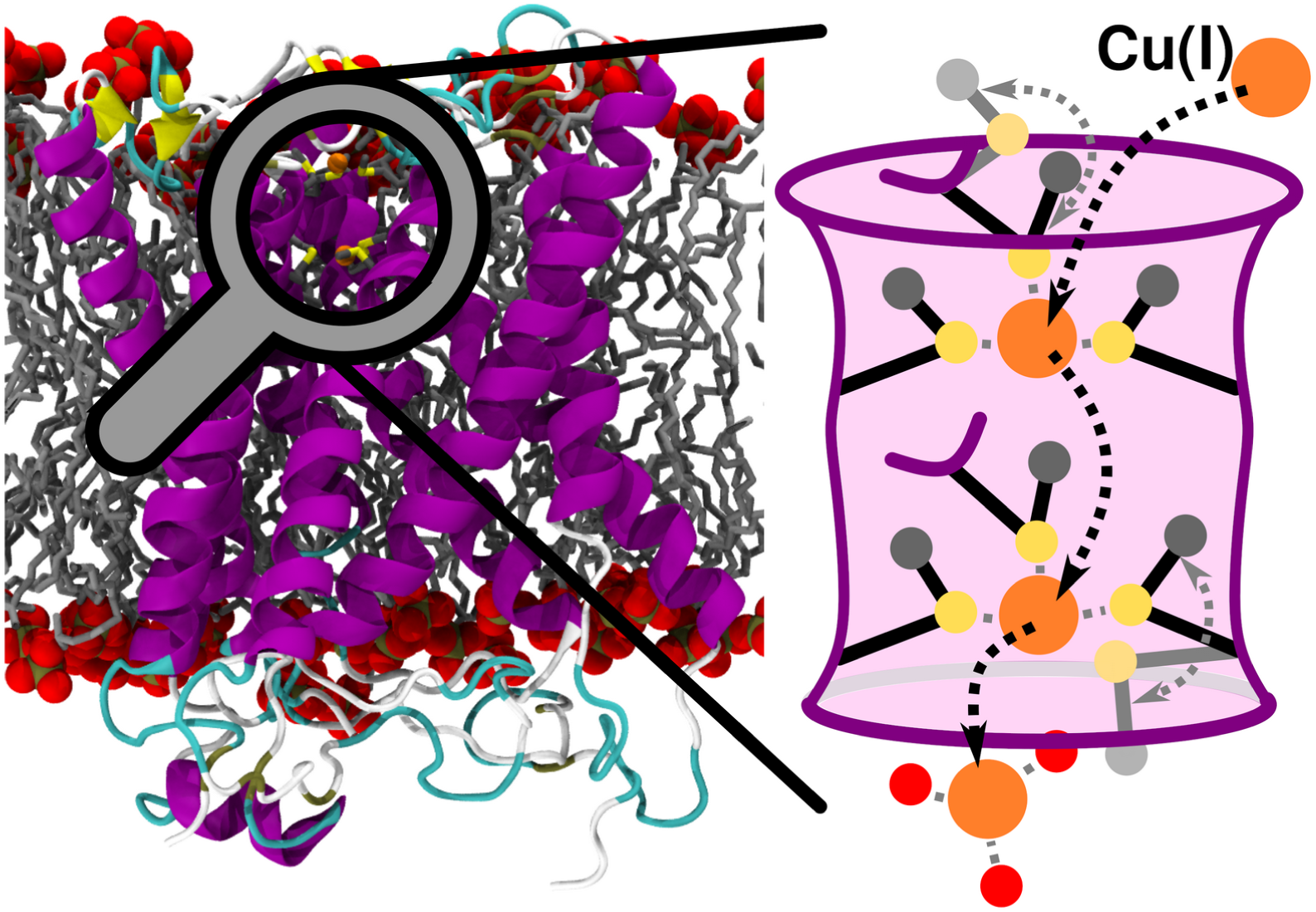

## 1. Introduction

Copper is an essential metal for cellular growth and development. Copper acts as a signaling agent, promotes electron or oxygen transport, and is a critical cofactor of copper-enzymes involved in a wide variety of biochemical processes, including aerobic respiration, superoxide detoxification, hormone and neuropeptide biogenesis and connective tissue maturation.^1^ As such, an imbalanced copper (Cu(I)) cellular uptake leads to metabolic abnormalities, anemia, neurological, cardiac, connective tissue, immune disorders and Menkes’ disease.^2^ Conversely, Cu(I) overload is linked to Wilson’s and neurodegenerative diseases (Prion’s, Parkinson’s, Alzheimer’s) and is overwhelmingly associated to cancer onset and progression.^3,4^

The transmembrane Copper Transporter 1 (CTR1) is an integral membrane protein mediating the selective Cu(I) uptake in all eukaryotes. CTR1, being the only known mammalian in-cell Cu(I) importer, plays a key regulatory role, tightly controlling the concentration of this trace metal to ensure proper cellular metabolism.^5^ Impaired CTR1-mediated Cu(I) transport causes delayed embryotic development, affects dietary Cu(I) acquisition, and alters cardiac and liver functions.^6^ As such, CTR1 is an appealing pharmacological target to tackle diseases linked to copper imbalance.^7^ Owing to its utmost importance CTR1 has been attracting the interest of bioinorganic chemists for decades, but mechanistic advances were hindered by the low resolution of the available structural data.^8,9^ Only the recent crystallographic structure from *Salmo salar*^6^ supplied a blueprint to unlock the mechanistic intricacies of the CTR1-mediated Cu(I) transport.

Human CTR1 is a 190-amino acids-long homotrimer composed of three parts: (i) The N-terminal domain is responsible for catching the Cu(II) from the extracellular space, its subsequent reduction to Cu(I) and transfer to the transmembrane (TM) part; (ii) The TM part, which shuttles the Cu(I) ions across the membrane; (iii) The intracellular C-term part, which delivers Cu(I) to the intracellular chaperones. While the N- and C-term are disordered and not (or only partially) resolved in the X-ray structure,^6^ the TM part is composed by three monomers, each arranged in a 3-helix bundle fold and bearing two methionine residues (M150 and M154, in human) in a MX3M motif. Altogether, the three MX3M motifs compose two sets of Methionine (Met) triads, one extracellularly- and one intracellularly-oriented, which form the CTR1 selectivity filter and can host up to two Cu(I) ions. Mutational studies showed that the highly conserved MX3M motif plays a fundamental role in the Cu(I) transport, since M150L and M154L substitutions in human CTR1 abolish copper influx.^10^ The role of these Mets is nevertheless ambiguous. On one hand, they most likely ensure the exclusive uptake of Cu(I) ions due to the copper’s chemical affinity for S-containing ligands; on the other hand, their bulky side chains can form a hydrophobic seal as observed in other channels.^6^

Here, by preforming enhanced sampling classical and quantum-classical (QM/MM) molecular dynamics (MD) simulations we unlock the molecular terms of the CTR1-mediated Cu(I) transport mechanism, disclosing an intriguing dual role of the selectivity filter Mets, which, owing to the conformational plasticity of their bulky side chains, strictly control the number and the rate of Cu(I) ions flowing through CTR1.

## 2. Methods

### 2.1 Model building

The structure of CTR1 comprising the transmembrane part and part of the C-term (residues 41-186) was taken from the crystallographic structure PDB entry: 6M98 from *Salmo salar* (Figure 1A).^6^ The chaperone apocytochrome b_562_RIL (BRIL), present in the crystal structure, was replaced by the native loop modelled with the MODELLER program.^11^ On the basis of this structure we built three models: one with only one Cu(I) ion bound to the most extracellularly exposed Met150 (M154 in hCTR1) triad (Site 1), one with a Cu(I) bound only to the intracellularly exposed Met146 (M150 in hCTR1) triad (Site 2) and one with two Cu(I) ions bound to Site 1 and Site 2 as observed in the crystal structure (Figure 1B).

**Figure 1.**
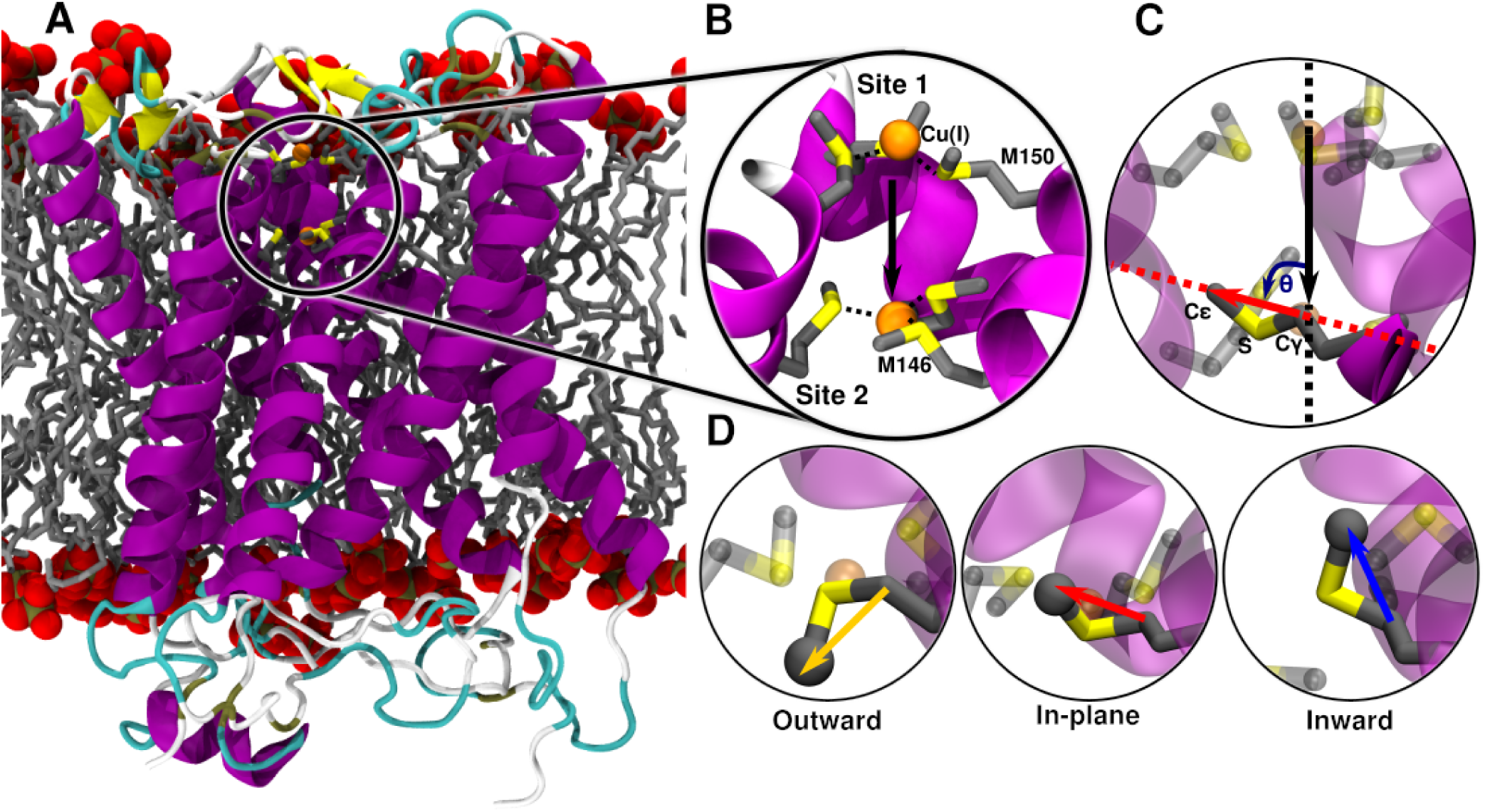
(A) Structure of CTR1 transporter from *Salmo salar* built on the X-ray structure (PDB code 6M98). CTR1 is shown in new cartoon representation, with magenta α-helices, yellow β-sheets and cyan loops. The membrane is represented as gray licorice, with the phosphate group displayed in van deer Waals spheres. (B) Close-up of the Met-triads with both Site 1 (up Methionine (Met)150 (M154 in hCTR1) triad) and Site 2 (bottom Met146 (M150 in hCTR1) triad) occupied with Cu(I) ions. Met residues are shown in licorice with S and C atoms in yellow and gray, respectively. Hydrogen atoms are omitted for clarity. Cu(I) is depicted as an orange van deer Waals sphere. Cu(I) coordination sphere is highlighted with black dashed lines. Black arrow indicates the vector the selectivity filter trace between centers of mass of the Site 1 and Site 2 Mets backbone atoms. (C) Definition of the θ angle used to classify the Met conformations: the black line indicates the vector of the selectivity filter and the red line indicates the Met Cγ-Cε vector. (D) The three possible conformations of the selectivity filter Site 2 Met146 with the Cε atom highlighted as gray sphere: outward, in-plane and inwards states highlighted with yellow, red and blue arrows, respectively.

CHARMM-GUI^12,13^ was used to insert the CTR1 protein into a lipid bilayer and create the simulation box. The orientation of CTR1 with respect to the membrane bilayer was determined using PPM webserver.^14^ The resulting membrane bilayer consisted of 95 phosphatidylcholine (POPC) molecules in each leaflet. The simulation box dimensions were 90×90×110 Å and further contained 17446 water molecules and 0.15 M concentration of NaCl (45 Na^+^ ions, 50 Cl^-^ ions). The protein was described using Amber FF14SB forcefield^15^ while for the POPC bilayer we used lipids17 FF.^16^ Water was described using the TIP3P model.^17^ Joung and Chetham parameters^18^ were used for Na^+^ and Cl^-^ ions. Cu FF parameters were taken from Merz and coworkers^19^ devised for MD simulations of Cys coordinating Cu-proteins. The system was minimized using 1000 steps of steep descent optimization with distance restraints on the S-Cu(I) coordination bonds with value 2.25 Å and force constant 5000 kJ/mol·nm. For the rest of the equilibration procedure the positions of Cu(I) ions and the methionine triads coordinating them were restrained using matrix of distance restraints with a force constant 2000 kJ/mol·nm. Gromacs 2018 was used for the classical MD simulations.^20^ The system was equilibrated in 8 steps with an increasing MD simulations length (0.5 ns, 0.25 ns, 0.5 ns, 0.5 ns, 0.5 ns, 0.5 ns, 0.5 ns, 2.0 ns) and decreasing restraints on the protein (5000 kJ/mol·nm, 4000 kJ/mol·nm, 3000 kJ/mol·nm, 2000 kJ/mol·nm, 1000 kJ/mol·nm, 500 kJ/mol·nm, 250 kJ/mol·nm, 50 kJ/mol·nm, 0 kJ/mol·nm) and the membrane (2000 kJ/mol·nm, 2000 kJ/mol·nm, 1000 kJ/mol·nm, 500 kJ/mol·nm, 250 kJ/mol·nm, 100 kJ/mol·nm, 50 kJ/mol·nm, 0 kJ/mol·nm, 0 kJ/mol·nm). Restraints on Cα atoms with 50 kJ/mol·nm were applied in the last step. The Cα restraints were kept for the first 30 ns of MD simulations to allow for membrane equilibration. 100 ns-long classical MD simulations of each model were performed before switching to QM/MM MD simulations.

### 2.2 QM/MM MD simulations

The QM/MM MD simulations were performed using CP2K version 6.1.^21^ The QM zone comprised all six Met residues (cut at the C - Cβ bond) forming the selectivity filter and one or two Cu(I) ions along with nearby water molecules. The QM zone of the system with the unoccupied bottom Met146-triad (used to study the Cu(I) translocation from Site 1 to Site 2) included 10 water molecules, which were close to Cu(I). These were periodically updated as the Cu(I) approached Site 2. Conversely, the QM zones of the simulations done to study the dissociation of Cu(I) from Site 2 towards the interval vestibule (comprising either two Cu(I) ions or only one Cu(I) in Site 2) included 8 waters molecules, which were restrained with an upper wall 6 Å distance restraint to the Site 2 Met146-triad defined by a geometric center of its Met C atoms. Additional restraint was placed on nearby MM water molecules with a lower wall distance of 6 Å to the Site 2 Met-triad to prevent the MM waters to exchange with the QM ones. The Plumed 2.7.0 plugin was used for the restraints.^22^

The QM zone was described using BLYP functional^23–25^ with dual Gaussian-type/plane waves basis set as in previous simulations.^26,27^ Specifically, a double-ζ (MOLOPT) basis set^28^ was used with an auxiliary PW basis set using the Goedecker–Teter–Hutter (GTH) pseudopotentials^29^ and a plane wave cutoff of 320 Ry. DFT-D3 dispersion correction was applied.^30^ After switching to QM/MM MD the system was relaxed through series of alternating short unbiased NVE and simulated annealing simulations using Langevin thermostat. Next, the system was gradually heated to 300 K over the course of 10 ps and left to equilibrate during a 10 ps unbiased QM/MM MD. The MD timestep was 0.5 fs. The heating and production runs used two canonical sampling through velocity rescaling (CSVR) thermostats:^31^ one applied to the QM part and the other to the rest of the system. Time constant of both was 100 fs.

### 2.3 QM/MM metadynamics simulations

The metadynamics (MTD) simulations were exploited to study the mechanism of Cu(I) translocation through the CTR1 selectivity filter. To this aim we employed the MTD implementation of CP2K using two collective variables (CVs): CV1 – distance of the Cu(I) to the plane of the Site 1 Met-150 triad, defined on the bases of the Met150 C atoms; CV2 defined as the coordination number (CN) of the Cu(I) with respect to Sulphur (S) atoms of the Site 2 Met146-triad similarly to a previous study.^32^ For the exponential form of coordination number we used the p=8, q=16 and d_0_=2.85 Å parameters. The width of the deposited MTD hills was 0.5 Å for CV1 and 0.05 for CV2, while the height was 1 kcal/mol and the deposition time was 50 fs. The effect of the MTD settings on the results was assessed by performing replicas MTD simulations with smaller hill height (0.6 kcal/mol) and longer deposition time (100 fs).

The dissociation of the Cu(I) ion from the Site 2 Met146-triad was simulated with the MTD implementation encompassed in Plumed 2.7.0 using again two CVs: CV1 – accounting for the z-projection of the distance of the Cu(I) to the center of the CTR1 selectivity filter defined using C atoms of all Site 1 and Site 2-Mets; and CV2 – defined as coordination number (CN) of the Cu(I) with respect to S atoms of the Site 2 Met146-triad. In this case the width of the deposited hills was 0.5 Å and 0.05, respectively for CV1 and 2, the height was 1 kcal/mol and the deposition time was 50 fs. The final dissociation step from the partially dissociated state, in which Cu(I) is still coordinated to one Site 2-Met, was explored with a separate set of MTD simulations using only one CV (CV1), accounting for the distance of the Cu(I) ion to the S atom of the last coordinated Site 2-Met. Restraints were put on the coordination number to the Site 2 Met146-triad at 1.15 value and the distance to the other Mets at 3.0 Å with a force constant 1000 kJ/mol·nm. The width of the deposited MTD hills was 0.2 Å, the height was 0.25 kcal/mol and the deposition time was 100 fs. When studying the Cu(I) dissociation from the CTR1 selectivity filter in a CTR1 model containing a single Cu(I) bound to Site 2, two replicas of the final MTD simulation were run to obtain more accurate free energy barrier. Conversely, when studying the same process for a CTR1 model hosting two Cu(I) ions, two replicas of the final dissociation MTD simulation were run, one with restraint on the χ_1_ and χ_3_ torsional angles of all Site 2-Mets to force them into the in-plane (IP)-conformation, and one unrestrained. This was done to explore the role of the water access to the Site 2 on the free energy barrier of Cu(I) in cell release.

### 2.4 Classical metadynamics simulations

Using the QM/MM MD-equilibrated structure we derived FF-parameters of the CTR1 Met Triads using the Metal Center Protein Builder (mcpb).^33^ Both Cu-occupied Met-triads were included in the “small model” used to parametrization the Cu(I) ion coordination sphere. The geometry optimization was done using Gaussian 09 at the B3LYP/6-31G* level with constraints on the distance between the Site 1 and Site 2 Mets. The resulting FF parameters were employed to equilibrate again the system using the previously described MD protocol and to explored the conformational behavior of the selectivity filter Met triads. Specifically, we performed MTD simulation to monitor the torsion angle behavior of the Mets. These can assume three different conformations within the selectivity filter: (i) Outward (O) with the S-Methyl bond parallel to the z-axis of the selectivity filter and the methyl pointing to away from the selectivity filter (i.e. towards the extracellular side for Site 1 and towards intracellular CTR1 vestibule for Site 2), (ii) In-Plane (IP) in which the S-Methyl bond is perpendicular to the z-axis of the filter and the Methyl group is pointing outside from the selectivity filter, and (iii) Inward (I) with the S-Methyl bond is parallel to the z-axis of the selectivity filter and with the methyl pointing inside the selectivity filter (Figure 1C).

In order to calculate the free energy cost for the conformational flipping between the IP and O state of one Site 2-Met the other two Site 2-Mets were blocked in their IP configuration by restraining the χ1 (_n-1_C-Cα-Cβ-Cγ) and χ3 (Cβ-Cγ-S-Cε) torsional angles to the range 2.77 – 3.14 rad. The conformational behavior of the third Site 2-Methionine was instead monitored by biasing χ1, χ2 (Cα-Cβ-Cγ-S) and χ3 torsion angles in a well-tempered MTD simulation using σ=π/18, height = 0.25 kcal/mol, bias factor 10 and deposition time 5 ps.

### 2.5 Analyses

For analyses purposes the three possible conformational states of Mets were defined, using the angle θ between two vectors: the first is the vector lying between geometric centers of Site 1 and Site 2 Met backbone atoms (Figure 1C), which represents the axis of the selectivity filter, the second vector is defined by Cγ-Cε Met atoms (Figure 1C). The Site 1 conformations are defined with θ angle ranges: 0-40° for the Outward (O), 50-90° for In-Plane (IP) and 100-160° for Inward (I) conformations of Met154 residues. The Site 2 conformations are defined with θ angle ranges: 100-160° for the Outward (O), 50-90° for In-Plane (IP) and 0-40° for Inward (I) conformations of the Met150 residues. The θ angle analysis along with that of the RMSD, RMSF, and radial distribution function was performed using cpptraj.^34^ The histograms of the θ angle distribution were created by numpy and density normalized.^35^ Met1-3 labels are used to refer to Mets coming from monomeric chain 1-3. In all MTD simulations the error of the free energy profiles was calculated as the standard deviation of different time averages from different simulation blocks. The minimum free energy paths were obtained using Metadynminer.^36^

## 3. Results

### 3.1 Binding mode of Cu(I) ions and conformational response of the selectivity filter

The crystal structure of CTR1 shows two Cu(I) ions bound to the selectivity filter, one per each Met-triad, with the Cu(I)-S distances in the upper (most-extracellularly exposed) Met150-triad (hereafter referred as Site 1) and in the bottom (most-cytosol exposed) exposed Met146-triad (hereafter refereed as Site 2) being 2.16 and 3.22 Å, respectively. The large distances between the Cu(I) ion and the sulfur atoms at Site 2 suggests that the metal ion is not (or is only weakly) bound to this site. The thioether group of the Met residues forming the triads supply a trigonal planar coordination geometry, presumably ensuring the selective uptake of Cu(I) over Cu(II) ions. Both Cu(I) ions are placed below the plane of the Met-triad to which they bind, laying at a distance of 7.22 Å from each other.

In order to refine the crystal structure and establish the number and the geometry of metal ions, which can simultaneously bind to CTR1, we performed a 100 ns-long classical MD equilibration followed by 10 ps QM/MM MD of the CTR1 model embedded in a lipid membrane mimic with two bound Cu(I) ions.^37^ The resulting average Cu(I)-S coordination distance for both triads is 2.32 ± 0.16 Å (Figure S1), while the Cu(I)-Cu(I) distance decreases to a mean value of 6.18 ± 0.42 Å (Figures 1B and S2). In order to assess whether the two Cu(I) ions may be spaced out by water molecules during their permeation, as in ions channels,^38^ we placed a water molecule between them. This invariably triggered a distortion of the selectivity filter (data not shown), leading us to rule out this possibility. As such, this set of simulations suggests that CTR1 may simultaneously bind two Cu(I) ions in the selectivity filter, even though the resulting coordination geometry is slightly different from that captured in the crystal structure (Figure S3). Additionally, we investigated the binding of only one Cu(I) ion to either the Site 1- or Site 2-Met triad. In this case, the Site 2 or Site 1-bound ion, respectively, was removed from the previous model, which was then subjected to additional QM/MM MD simulations. The coordination geometries obtained in the presence of a single Cu(I) ion were similar to those of the two Cu(I) ions-bound model, but a remarkable difference in the conformational plasticity of the selectivity filter Mets emerged. First, the RMSD and RMSF analyses of these residues pinpointed striking asymmetries between the two Met-triads (Figures S4-S5) in the distinct models. Namely, while the overall flexibility of Site 1 Met-triad increases when CTR1 hosts two Cu(I) ions in the selectivity filter as compared to the Site 1 only-bound model, the flexibility of one of the Site 2 Mets layer drastically decreases in the presence of two Cu(I) ions with respect to the Site 2 only-bound model (Figure S5).

A detailed inspection of the conformational behavior of the selectivity filter Mets revealed that these residues can assume three different conformations (Figure 1D): (i) An Outward (O)-state, where the S-methyl bond is parallel to the z-axis of the selectivity filter and the methyl points far away from the selectivity filter center (i.e. it points towards the extracellular side in Site 1 and towards intracellular CTR1 vestibule in Site 2), (ii) an In-Plane (IP)-state, in which the S-methyl bond is perpendicular to the z-axis of the filter and the Methyl group points away from the selectivity filter center, and (iii) an Inward (I)-state with the S-methyl bond parallel to the z-axis of the selectivity filter and with the methyl group pointing inside the selectivity filter.

The analysis of Mets conformational behavior (Figure 2) disclosed clear differences among the distinct models investigated. In the apo model the Mets residues access all conformations, even though a coupling exists between the Site 1 and Site 2 Met-triads with only one Site 1-Met at time filling the selectivity filter by assuming the I-conformation. When only Site 1 is occupied two coordinating Mets assume IP-conformation and one lies in between the IP- and O-state. Conversely, the Mets of the empty Site 2 are more flexible with one laying in the IP-state, one between the IP and I-state and one toggling around the IP-conformation reaching at times the I- or O-state (Figure 2). Interestingly, when only Site 2 is occupied, two Mets coordinating the Cu(I) ion retain the IP-conformation, with one Met still toggling around the IP-state, and visiting more frequently the O-conformation. Conversely, two Mets of the empty Site 1 assume the IP-conformation and one the O-state (Figure 2). The latter Met most likely hinders the access of additional Cu(I) ions to the selectivity filter from the extracellular matrix. Stunningly, when both sites bind a Cu(I) ion, all Site 1-Mets rigidly retain the IP-conformation, while one of the Site 2-Mets lays between the I- and IP-conformation, one in the IP conformation, and the last one remains rigidly trapped in the O-conformation. This reveals that the conformations assumed by of the two Met-triads are not only coupled, but also intimately entwined with the number of bound Cu(I) ions. As such, their Cu(I)-induced conformational selection may play a key role in the CTR1 transport mechanism. To further elucidate this intriguing trait, in the following, we investigated the Cu(I) transport mechanism through CTR1 in the presence of one or two Cu(I) ions bound to the selectivity filter.

**Figure 2.**
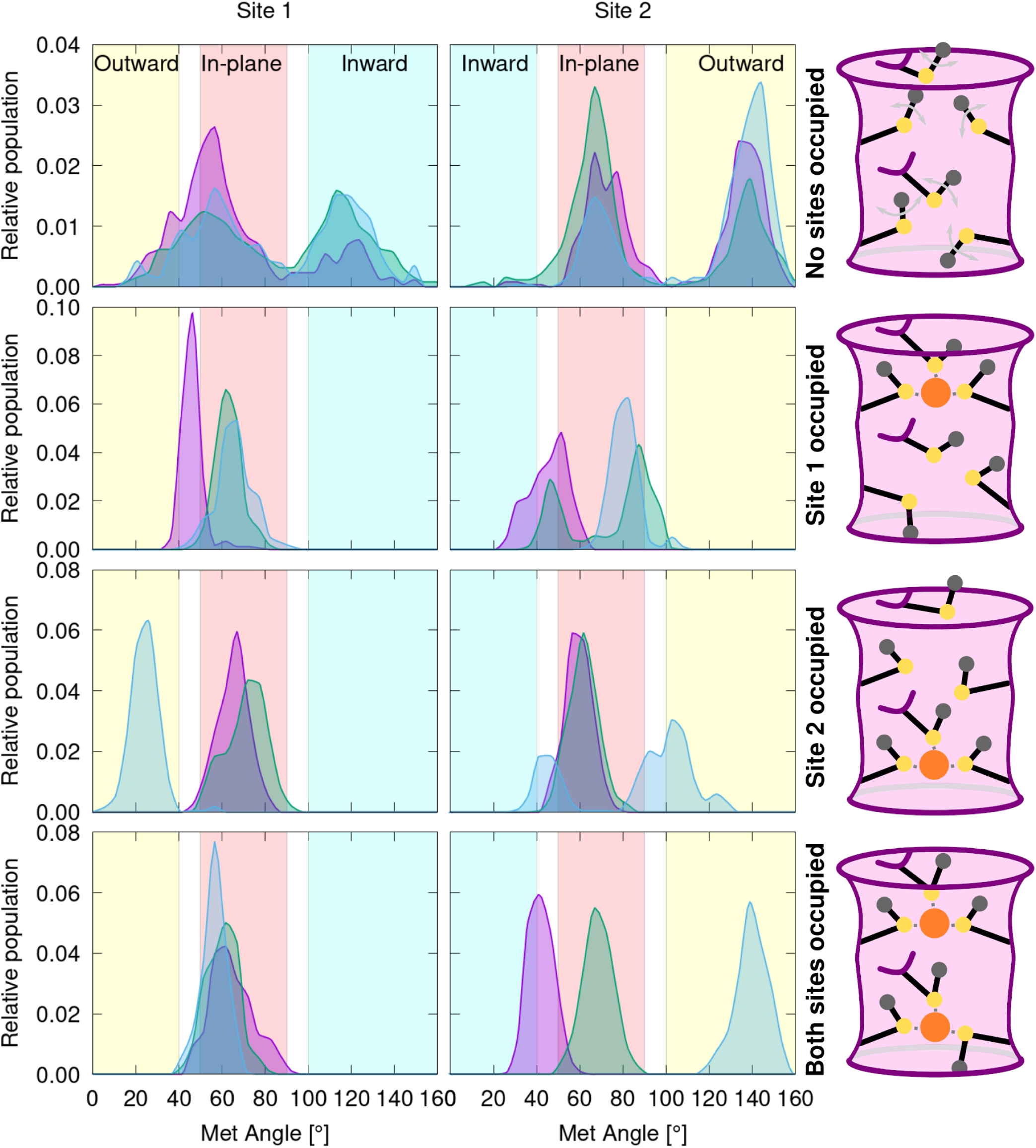
Distribution of the angle (°) (defined as in Figure 1C) as obtained from QM/MM and classical (for apo CTR1, containing no bound Cu(I) ions) molecular dynamics trajectories. The distribution is shown for apo CTR1 model (top panel), a Cu(I) ion bound to Site 1 (second panel from top); to Site 2 (third panel), and with Cu(I) ions bound to both sites (bottom panel). Left and right column show the distribution of the Site 1 and Site 2 Met-triads, respectively. The distributions of the three Met1-3 residues are shown with different colors (magenta, green and blue, respectively). Areas of the histogram corresponding to the inward, in-plane and outward-conformations are highlighted in cyan, red and yellow, respectively. Sketches of the conformational behavior of the models are shown on the right. The CTR1 selectivity filter is schematically depicted as a pink cylinder with violet walls. The Mets residues are schematically represented in black lines with Sulphur, Cu(I) and Cε atoms highlighted in yellow, orange and gray circles, respectively. Cu(I) coordination sphere is shown with gray dashed lines. In apo CTR1 gray arrows indicate the Met conformational motion.

### 3.2 Cu(I) translocation from the Site 1 to the Site 2 Met-triad

We initially assessed the early events of CTR1-mediated transport by monitoring the translocation of a Cu(I) ion from Site 1 to Site 2. CTR1 model initially hosted Cu(I) in Site 1 and the CTR1 import mechanism was investigated by performing QM/MM MTD simulations using two collective variables (CVs). The first (CV1) was defined as the distance of the Cu(I) ion from the plane of the Site 1-Met triad and the second (CV2) as the coordination number (CN) of the Cu(I) ion with respect to the S-atoms of the Site 2-Met triad (Figure 3). These MTD simulations unveiled that the Cu(I) ion, by surmounting a Helmholtz free energy barrier (ΔF^‡^) of 5.8 ± 2.8 kcal/mol, heads from Minimum 0 (M0), where it is coordinated by the three Site 1-Mets, all in the IP-state, to M1, lying at a free energy (ΔF) of −4.9 ± 2.8 kcal/mol, where Cu(I) is bound by two Site 1-Mets, in the IP-state, and one Site 2-Met also in the IP-state. Next, Cu(I) moves to M2 (ΔF^‡^ = 6.1 ± 1.1 kcal/mol and ΔF =-3.9 ± 0.2 kcal/mol), where it is bound by one Site 1-Met in the IP-state and two Site 2-Mets in the IP-state. Interestingly, the Site 1-Met1, still coordinating the metal, moves down within the selectivity filter to follow Cu(I), while maintaining the IP-conformation (Figure S6C,E). The Cu(I) ion ultimately reaches M3 (ΔF^‡^ = 4.3 ± 4.0 kcal/mol), where it is coordinated by the three Site 2-Mets, all in the IP-state (Figure S6C,D).

**Figure 3.**
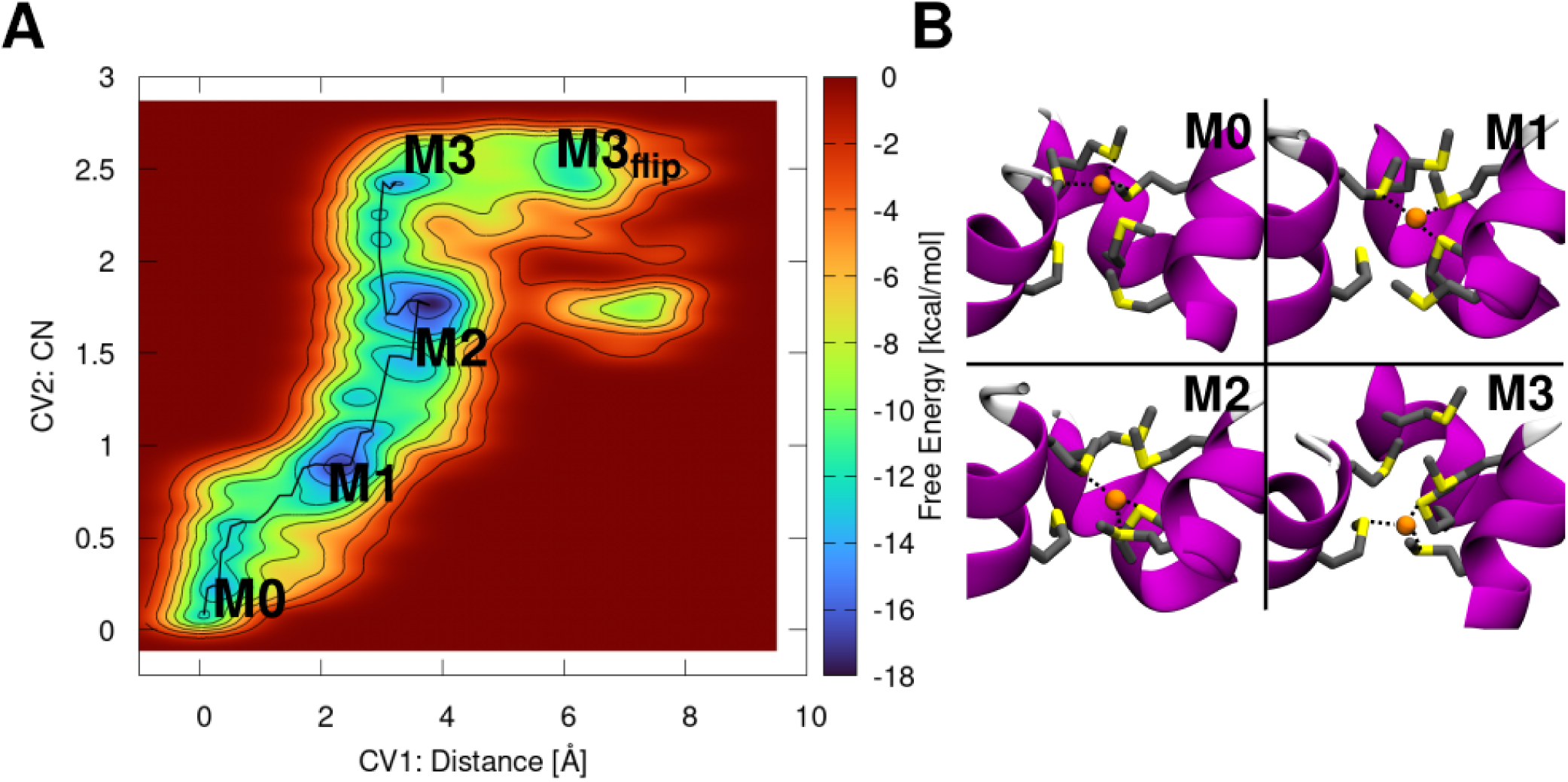
(A) Free energy surface (FES, kcal/mol) of the Cu(I) translocation from Site 1 to Site 2 plotted as function of Cu(I) distance from the Site 1 Met150-triad (CV1) and coordination number of Cu(I) with respect to the bottom Site 2 Met154-triad (CV2). The FES is shown from blue to red with isosurface lines drawn every 2.0 kcal/mol. (B) Close-ups of the minima visited during the translocation. CTR1 is shown as pink new cartoon, Met-triads are displayed in licorice with Cu(I) ion shown as an orange sphere. Hydrogens are omitted for clarity. The Cu(I) coordination sphere is highlighted with gray dashed lines.

The additional minimum observed in the FES (M3_flip,_ Figure S6) is associated to the conformational switch of one Site 2-Met from IP to O-conformation, which occurs after Cu(I) is fully transferred to the Site 2 (Figure S6D,F). A ΔF^‡^ = 5.9 ± 1.0 kcal/mol separates M3 from M3_flip_. Even if in M3_flip_ the Cu(I) ion has moved further down towards the cytosolic exit as compared to M3, suggesting that M3_flip_ might be an early intermediate along the Cu(I) dissociation pathway from Site 2 to the cytosol, the O-conformation of the Site 2-Met may prevent the access of water to the selectivity filter and thus the Cu(I) dissociation towards the cytosol-exposed cavity of CTR1. As such, in the following, we investigated the Cu(I) dissociation from the selectivity filter starting from M3, which is more thermodynamically stable than M3_flip_ and displays all Site 2-Mets in the IP-state, allows for the water molecules to easily access the Cu(I) coordination sphere (Figure S7), thus facilitating the Met/water exchange reaction and possibly the Cu(I) dissociation towards the cytosolic cavity.

### 3.3 In-cell Cu(I) dissociation mechanism

From M3 state, two possible scenarios open: the Cu(I) ion can dissociate from the selectivity filter to the cytosol before or after the binding of a second Cu(I) ion to the Site 1.

#### 3.3.1 Cu(I) dissociation from Site 2 in the absence a Cu(I) ion bound to Site 1

We initially assumed that a single Cu(I) ion at time flows through CTR1 and explored the dissociation of Cu(I) from Site 2 in the absence of any Cu(I) ion bound to Site 1. To this aim we performed QM/MM MTD simulations using as CVs the z-projection of the Cu(I) distance from the center of CTR1 selectivity filter (CV1) and the coordination number (CN) of Cu(I) with respect to the Site 2 Met-triad Sulphur atoms (CV2) (Figure 4). This MTD simulation was started from the equilibrated structure of the M3 state (Figure 3). From M3 the Cu(I) ion in-cell release starts by overcoming a ΔF^‡^ = 5.8 ± 2.6 kcal/mol barrier to reach the M4 state (ΔF= −3.0 ± 2.7 kcal/mol), where Cu(I) acquires a water molecule in the coordination sphere, after losing one Site 2-Met. Next, Cu(I) reaches M5 (ΔF^‡^ of 8.5 ± 2.7 kcal/mol, ΔF = 3.0 ± 3.2 kcal/mol), where it is hydrated by two water molecules, while being still bound to a single Site 2-Met. Finally, the Cu(I) ion reaches M6, in which it is coordinated by three waters (ΔF^‡^ = 12.5 ± 1.2 kcal/mol (Figure S9),^a^ ΔF = 8.1 ± 3.0 kcal/mol). The Site 2-Met1, still coordinating the Cu(I) ion in M5, is very flexible, toggling between IP-, I- and O-conformations during the dissociation process (Figure S8), while the other two Met residues remain in IP-state (M4-M6 states). This highlights how the Cu(I)-dependent conformational remodeling of the selectivity filter Mets takes an active role during Cu(I) transport.

**Figure 4.**
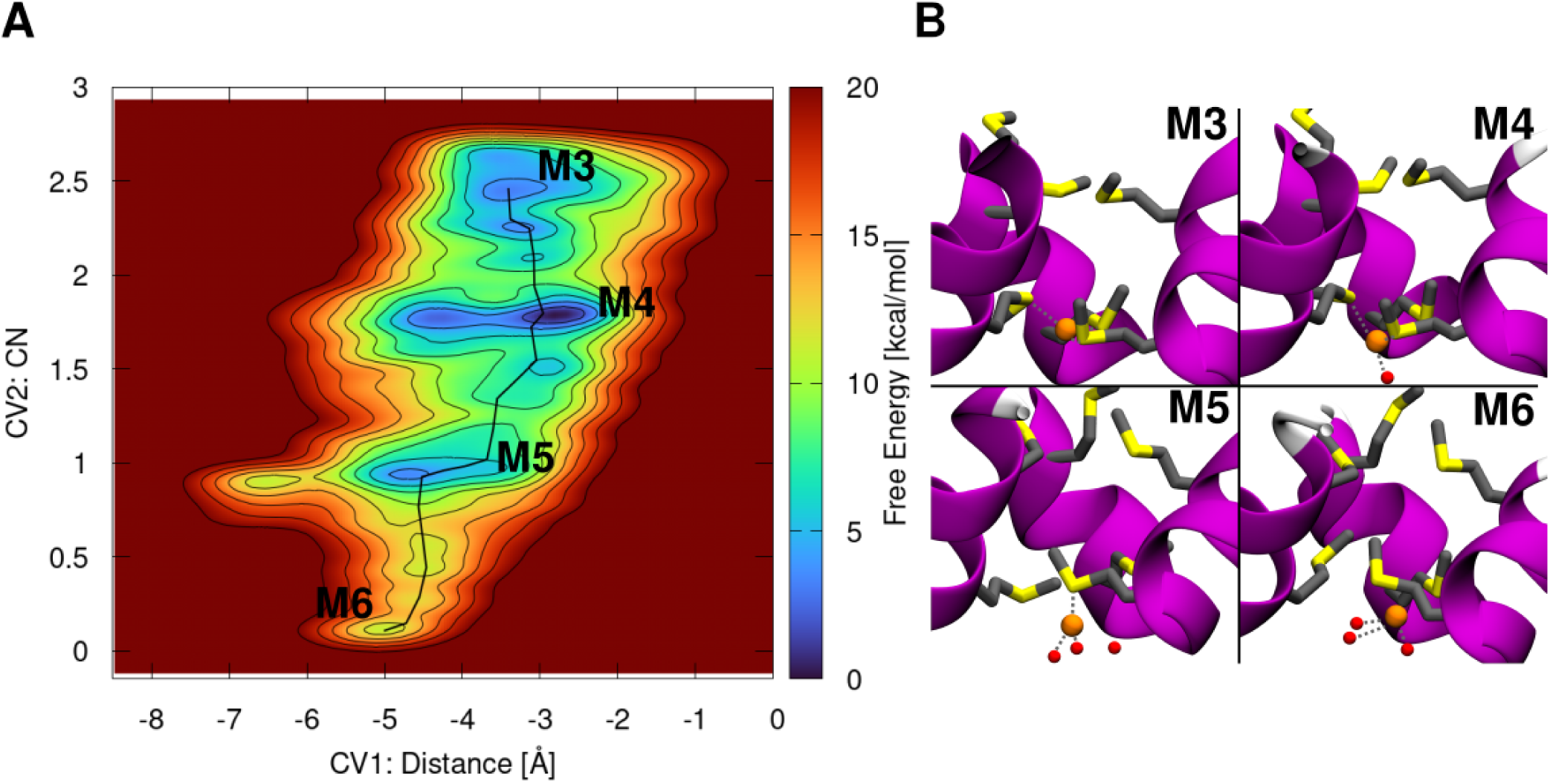
Mechanism of the Cu(I) dissociation from Site 2 in absence of Cu(I) in Site 1. (A) Free energy surface (FES, kcal/mol) plotted as a function of the z-projection of the Cu(I) distance (Å) from the center of the selectivity filter defined by Cα atoms of the Site 1 and Site 2 Mets (Collective Variable 1, CV1); and coordination number (CN) of Cu(I) with respect to the Site 2 Met-triad (CV2). The FES is shown from blue to red with isosurface lines drawn every 2.0 kcal/mol. (B) Close-up views of Cu(I) dissociation states from Site 2. CTR1 is shown as magenta new cartoons, Met-triads as licorice, Cu(I) ion and water molecules as orange and red spheres, respectively. Hydrogens are omitted for clarity. Cu(I) coordination is highlighted with gray dashed lines.

#### 3.3.2 Cu(I) dissociation from Site 2 in the presence of Cu(I) ion bound to Site 1

We finally explored the dissociation of Cu(I) from Site 2, when also Site 1 binds a Cu(I) ion, by using the same set of CVs as detailed above. In the initial M_2Cu_0 state, the two Cu(I) ions force one Site 2-Met to assume a conformation in-between the IP- and I-states, while the remaining two Mets adopt the IP- and the O-conformation (referred as Met1, Met2 and Met3, respectively). Next, the metal dissociates from Site 2 by initially losing the coordination with the Site 2-Met1 in the I-state (M_2Cu_1; ΔF^‡^ = 10.8 ± 2.7 and ΔF = 8.9 ± 2.4 kcal/mol). This is followed by a water exchange reaction with the Site 2-Met3 in the O-state reaching the M_2Cu_2 intermediate (ΔF^‡^ 18.9 ± 3.1 kcal/mol, ΔF = 17.8 ± 2.9 kcal/mol, Figure 5). Noticeably, in M_2Cu_2 the Cu(I)-Cu(I) distance as well as the distances between Cu(I) and the Site 2-Met residues resembles those captured in the X-ray structure.^6^

**Figure 5.**
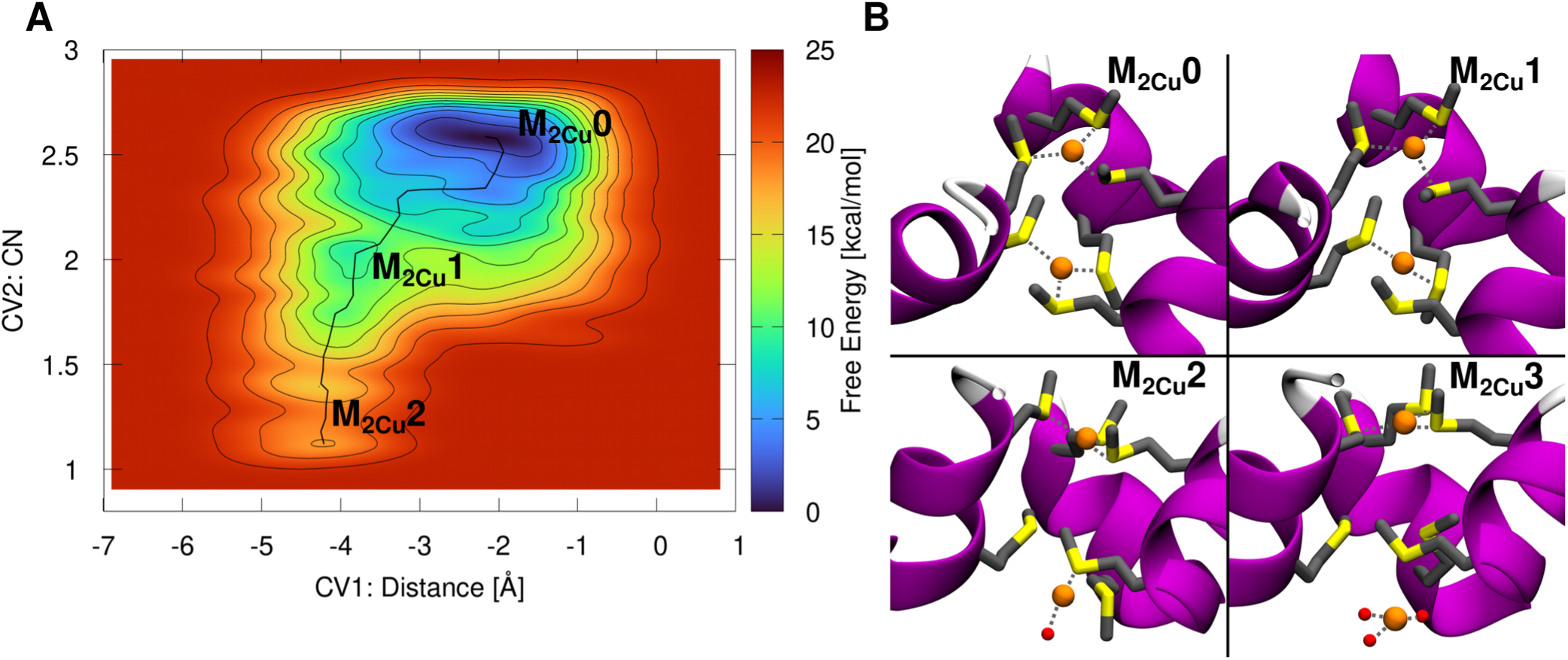
Mechanism of the Cu(I) dissociation from Site 2 in presence of Cu(I) in Site 1. (A) Free energy surface (FES, kcal/mol) plotted as function of the z-projection of the Cu(I) distance (Å) from the center of the selectivity filter defined by Cα atoms of the Site 1 and Site 2-Mets (Collective Variable 1, CV1); and coordination number (CN) of Cu(I) with respect to the Site 2 Met-triad Sulphur atoms (CV2). The FES is shown from blue to red with isosurface lines drawn every 2.0 kcal/mol. (B) Close-ups of the states visited during Cu(I) dissociation from Site 2 (state M_2Cu_3 obtained from additional 1D MTD, Figure S12). CTR1 is shown as magenta new cartoon, Met-triads as licorice and Cu(I) ion and water molecules as orange and red spheres. Hydrogens are omitted for clarity. Cu(I) coordination is shown with gray dashed lines.

Stunningly, in this intermediate-state the Site 2-Met2, which is the last Met coordinating Cu(I), is rather flexible, approaching the O-state during the final dissociation step (Figure S10), while Site 2-Met3, which permanently adopts the O-conformation, occludes the water access to the selectivity filter, thus hampering the Met/water exchange reaction (Figure S7) and hindering the complete dissociation of Cu(I) from Site 2. To assess how the Site 2 hydration affects the energetics of the Cu(I) in-cell release we performed additional MTD simulations using as CV the distance of Cu(I) to the Site 2-Met2. To this aim we built two different models based on M_2Cu_2 intermediate: one with the Site 2-Mets free to move (i.e. initially assuming one the O-, and two the IP-state); and a second model in which all Site 2-Mets were restrained to the IP-conformation (Figure S12). The resulting ΔF^‡^s to reach the fully dissociated M_2Cu_3 state were of 32.2 ± 2.7 and 21.3 ± 3.3 kcal/mol for the two models, respectively (Figure S12), confirming that the Met3 in the O-state hinders Cu(I) in-cell release. Nevertheless, the reduced conformational plasticity of the Mets, induced by restraining them in the IP-conformation, also contributes to slowing the Cu(I) in-cell release (increasing the ΔF^‡^ to 21.3 ± 3.3 kcal/mol) as compared to the Cu(I) dissociation from the Site 2-only bound model where all Mets were in the IP-state without any constraints (ΔF^‡^ = 12.5 ± 1.2 kcal/mol, Figure 4). Additionally, we monitored the Gibbs free energy cost (ΔG^‡^) of flipping a single Site 2-Met from the O-to IP-state, in the two Cu(I) ions-bound model, by performing classical MTD simulation using the dihedral angles χ1, χ2 and χ3 as CVs (Figure S13). The resulting ΔG^‡^ was of 8.6 ± 1.9 kcal/mol, further indicating that the Met conformation flipping occurs at a non-negligible free energy cost.

These findings compellingly show that the Cu(I) binding to both Met-triads reciprocally affects their conformational plasticity, with the Mets side chains acting as gate to limit the amount of in-cell Cu(I) import.

## 4. Discussion

Our atomic-level dissection of the CTR1-mediated Cu(I) in-cell uptake mechanism discloses that during the early steps of Cu(I) translocation from the Site 1 to Site 2, the metal ion visits two stable intermediates in which Cu(I) is concomitantly coordinated to both Site 1 and Site 2-Mets. The progression between these two minima displays the highest ΔF^‡^ (6.1 ± 1.1 kcal/mol) for Cu(I) movement within the selectivity filter (Figure 6). Nevertheless, the rate-limiting step of the in-cell transport process corresponds to the Cu(I) dissociation from the selectivity filter towards the cytosol-exposed CTR1 vestibule. The calculated ΔF^‡^ = 12.5 ± 1.2 kcal/mol (Figure 6) of this step is in good agreement with experimental turnover rate of 6.0-14.2 s^-1^ at 37 °C^39^ corresponding to a ΔG^‡^ of 13.0-13.5 kcal/mol.

**Figure 6.**
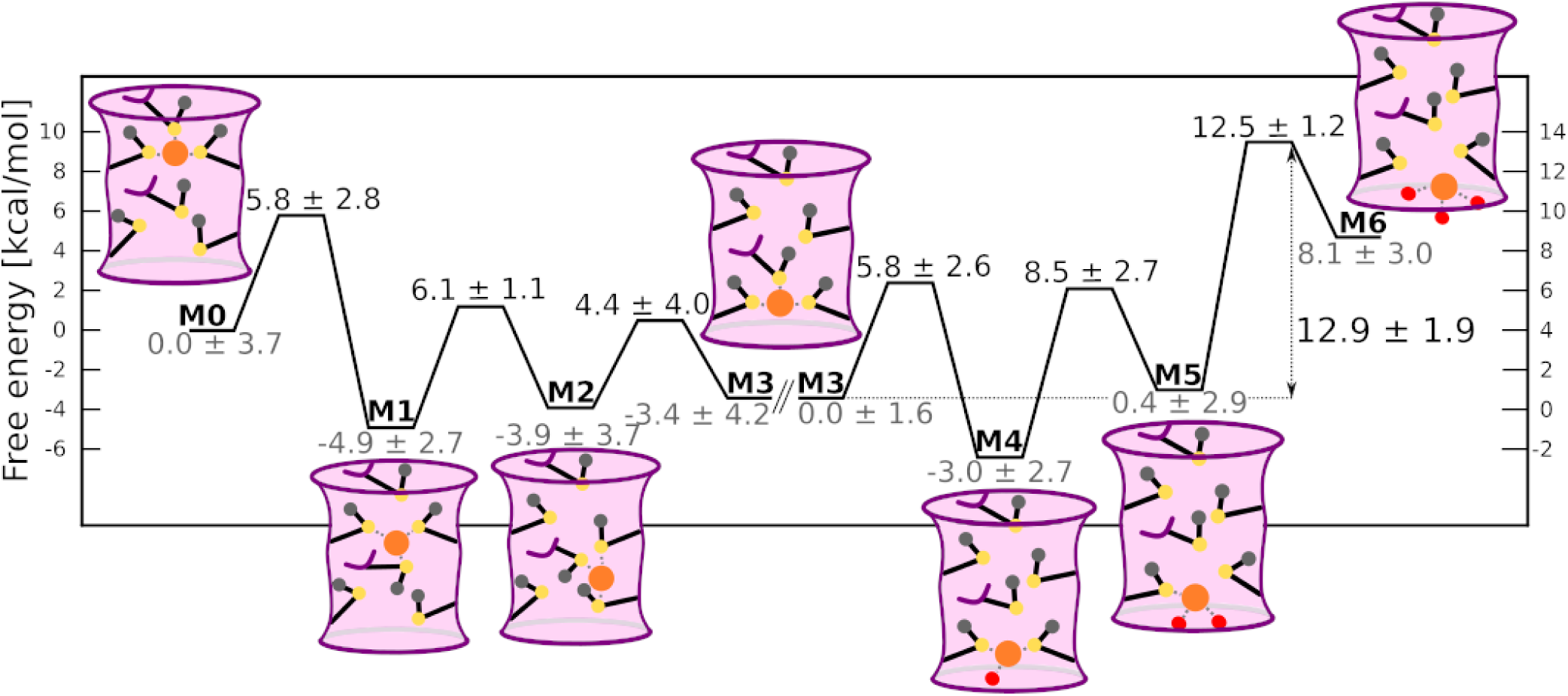
Schematic representation of the CTR1-mediated Cu(I) translocation processes. Free energy barriers of each step are reported (kcal/mol). The CTR1 selectivity filter is schematically depicted as a pink cylinder with violet walls. The Mets residues are schematically represented in black lines with Sulphur, Cu(I) and water oxygen atoms highlighted with yellow, orange and red circles, respectively. Cu(I) coordination sphere is shown with gray dashed lines. Parallel diagonal lines separate the results of the Cu(I) translocation within the selectivity filter from those of Cu(I) release to the cytosol. The left and right axes refer to the free energy cost for translocation and dissociation mechanism, respectively.

Most importantly, our findings unprecedentedly unlock an intimate coupling between the Site 1 and Site 2-Met triad occupancy and conformational plasticity as a salient CTR1 trait. Owing to this coupling, one Site 2-Met adopts an O-conformation when two Cu(I) ions are bound to CTR1, which prevents Cu(I) hydration (i.e. water access to the Cu(I) bound at the intracellular exit of the selectivity filter), significantly increasing the free energy barrier (ΔF^‡^ to 32.2 ± 2.7 kcal/mol) for the Cu(I) in-cell release as compared to the same step in the Site 2 only-bound model (ΔF ^‡^ = 12.5 ± 1.2 kcal/mol). Altogether, our outcomes disclose that CTR1 sophisticatedly leverages the conformational coupling of the Met-layers in the selectivity filter to selectively propel Cu(I) in-cell uptake in a limited and healthy amount (i.e. enabling the translocation of a single Cu(I) ion at the time). In this scenario, it is tempting to argue that when two Cu(I)-ions bind to CTR1, as it may potentially occur at high extracellular copper concentrations, the conformational coupling of the two selectivity Met-triads may be an elegant way of limiting the Cu(I) import rate. This may thus represent a sophisticated autoregulatory mechanism to protect cells from excessively high, and hence toxic, in-cell Cu(I) levels.

The importance of dynamic properties of coordinating residues, dictated by local interactions, for selective ion transport has been highlighted before, notably for the potassium channel KcsA.^40^ Similar to CTR1, KcsA contains multiple metal binding sites, however binding of multiple potassium ions is thought to ensure highly efficient ion conduction via so called knock-on mechanism mediated by electrostatic repulsion between potassium ions.^41^ Contrary, we find binding of multiple Cu(I) ions in the selectivity filter of CTR1 decreases the copper influx rate, most likely reflecting the different roles potassium and copper ions play in the cellular milieu.

Mets residues were shown to participate in ion channel gating previously, albeit as a hydrophobic barrier, precluding the ion passage indirectly,^42,43^ not through a clever inter-play between metal coordination and their conformational plasticity as in CTR1. Interestingly, Met-mediated Cu(I) transport is shared also by the bacterial CusA efflux pump relying on three sets of Met-diad binding sites to rapidly export Cu(I). However, these Mets do not seem to play a regulatory role, most likely due to the bacterial need to rapidly expel toxic Cu(I) ions from the cells to guarantee their survival.^44^ This confirm the hypothesis that the third Met in each binding site may be necessary to achieve autoregulatory function.

## Conclusion

In this study QM/MM MD enhanced sampling simulations resolve the intricate mechanism underlying the Cu(I) import through CTR1, compellingly assigning a new dual functional role to the selectivity filter Met residues. These residues selectively bind Cu(I) ions thanks to the affinity of their sulphur atoms for copper and also act as gates to regulate Cu(I) import uptake in a controlled and healthy amount.^6^ Namely, our outcomes show that the selectivity filter Mets act as gates in response to Cu(I) overload thanks to a sophisticated coupling between the amount of Cu(I) ions bound to CTR1 and the conformational response of the Met side chains. Indeed, .the simultaneous binding of two Cu(I) ions to the selectivity filter activates the Met gate, reducing the Cu(I) ions flow-rate to avoid their imbalanced, and hence toxic, accumulation in the cell. The detailed atomic-level picture of CTR1 import mechanism supplied here sets a conceptual basis to develop novel mechanism-based therapeutics tackling the variety of human diseases entwined to an inappropriate Cu uptake.

## Supporting information

Movie M1: Cu(I) translocation from Site 1 to Site 2

Movie M2: Cu(I) dissociation from Site 2

## Authors contributions

A.M. designed and supervised the study. P.J. performed the simulations. P.J. and J.A. analyzed the results. P.J., J.A., A.M. and S.R. wrote the manuscript. A.M. and S.R. secured the funding.

## Data availability

Trajectory data for all simulations can be obtained from authors upon request.

## SUPPORTING INFORMATION

The Supporting Information is available free of charge at https://pubs.acs.org/.

These include Figures S1 to S13: Cu(I)-S and Cu(I)-Cu(I) distances, Met-triads structure, RMSD and RMSF of Site 1 and Site2, Cu(I) translocation from Site 1 to Site 2, RDF of Cu(I)-Owater distances, Cu(I) dissociation from the Site 2 in absence of second Cu(I), Cu(I) dissociation from the Site 2 in presence of Site 1 Cu(I), O-to IP-conformational flipping from classical MTD; and Movies: M1 - Site 1 to Site 2 Cu(I) translocation and M2 - Site 2 Cu(I) dissociation.

## Acknowledgments

PJ was supported by the HPC Europa Fellowship (HPC17M882U) under the Project HPC-EUROPA3 (INFRAIA-2016-1-730897). AM thanks the financial support of Italian Association for cancer research (AIRC) project Investigator Grant #24514 and the ‘Against bRain cancEr: finding personalized therapies with in Silico and in vitro strategies’ (ARES) CUP: D93D19000020007 POR FESR 2014 2020 - 1.3.b - Friuli Venezia Giulia for financial support. JA was supported by the ‘Giovanni Fraviga’ AIRC-FIRC fellowship. The authors thank Exact Lab for the computational resources available through the ARES project.

## Footnotes

a The free energy barrier obtained from the original metadynamics simulations based on two collective variables is of 9.8 ± 3.5 kcal/mol.

