## Supporting Figures S1-S13 for "The Conformational Plasticity of the Selectivity Filter Methionines Controls the In-Cell Cu(I) Uptake through the CTR1 transporter"

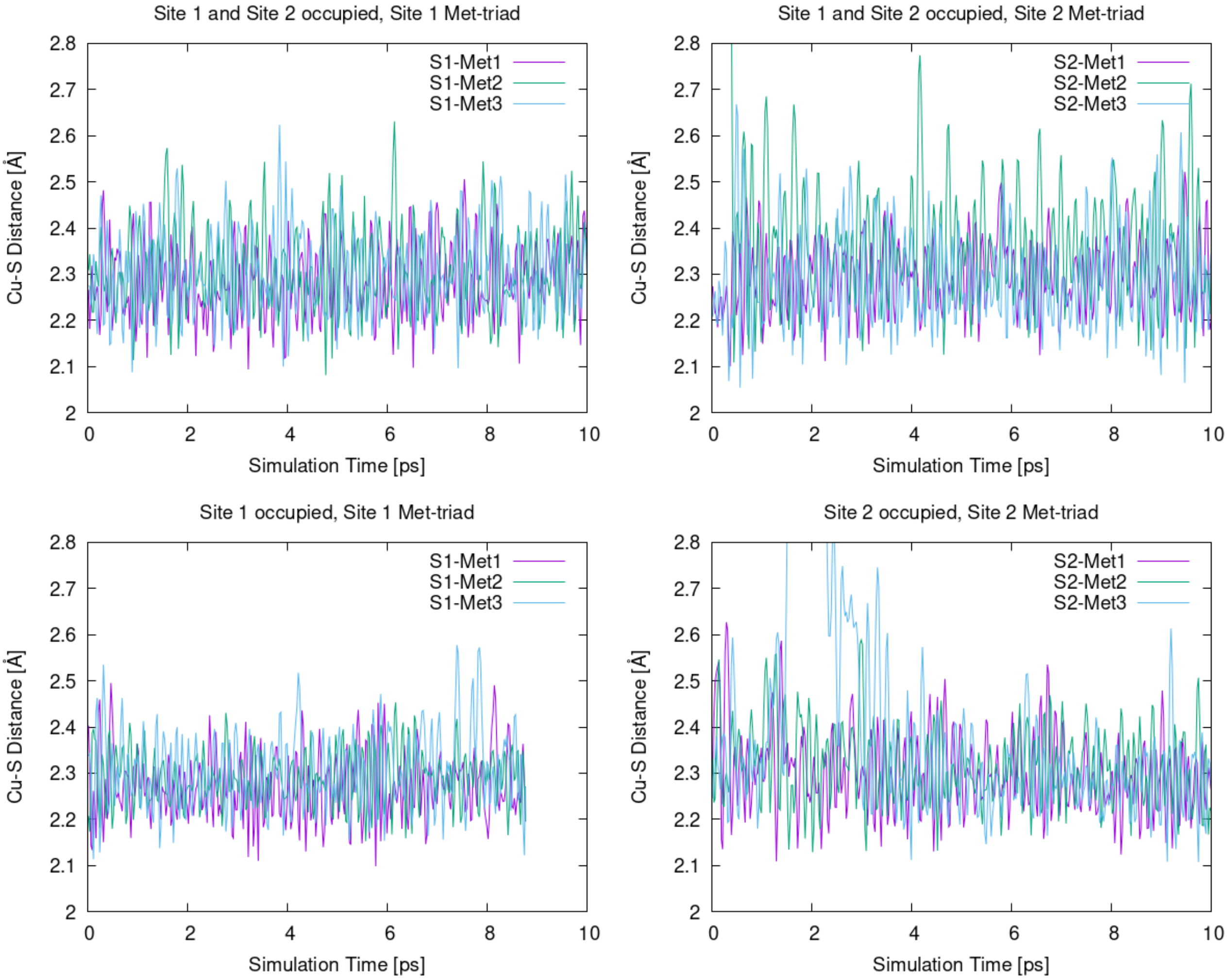

**Figure S1.** Cu(I)-S distances (Å) vs QM/MM molecular dynamics simulation time (ps) from the methionine residues of the CTR1 selectivity in Site 1 (top or most extracellular oriented) and Site 2 (bottom, most cytosol oriented) triad with both triad binding a Cu(I) ion (A and B), with a Cu(I) bound only to Site 2 (C) and only to Site 1 (D).

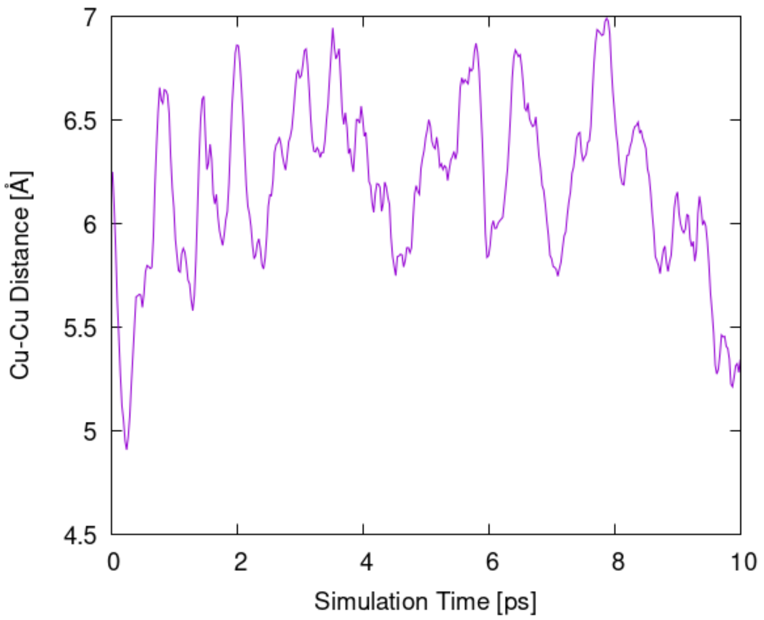

**Figure S2.** Cu(I)-Cu(I) (Å) distance vs QM/MM molecular dynamics simulation time (ps) in the CTR1 model with both Met-triads occupied by Cu(I) ions.

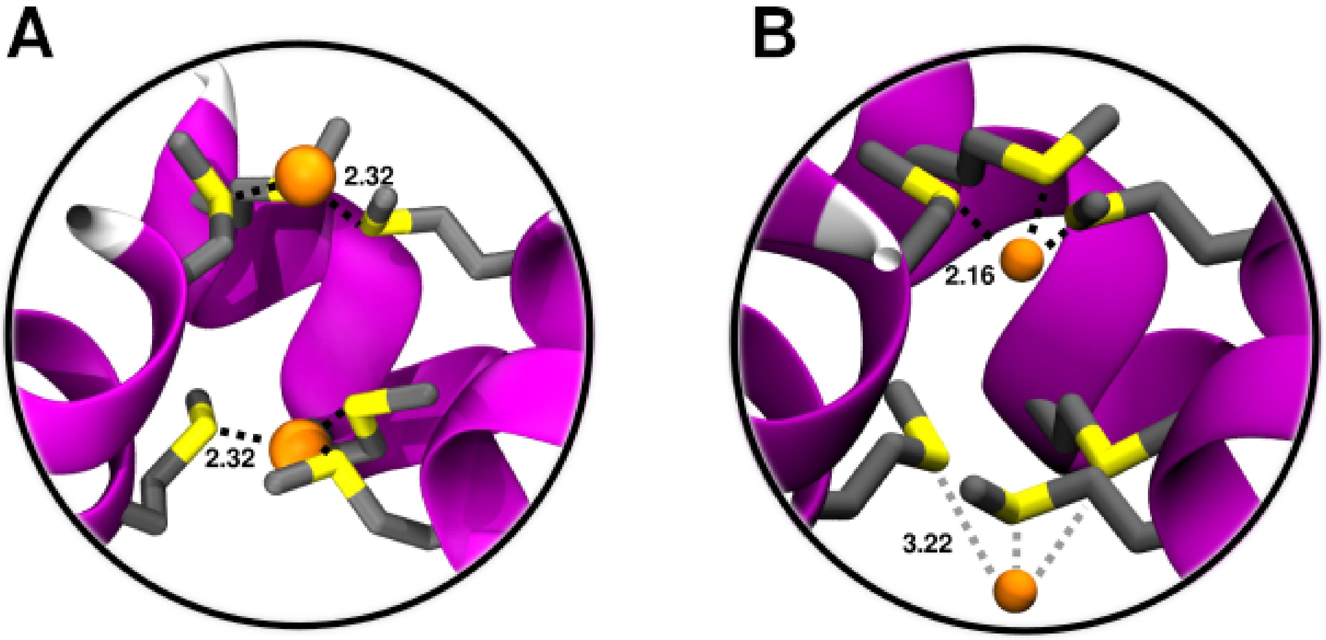

**Figure S3.** A) Structure of the Site 1 (upper) and Site 2 (lower) Met-triads each coordinating a Cu(I) ion as obtained from QM/MM molecular dynamics simulation. B) Structure of the Met-triads each binding a Cu(I) as obtained from the X-ray structure (PDB ID: 6m98).

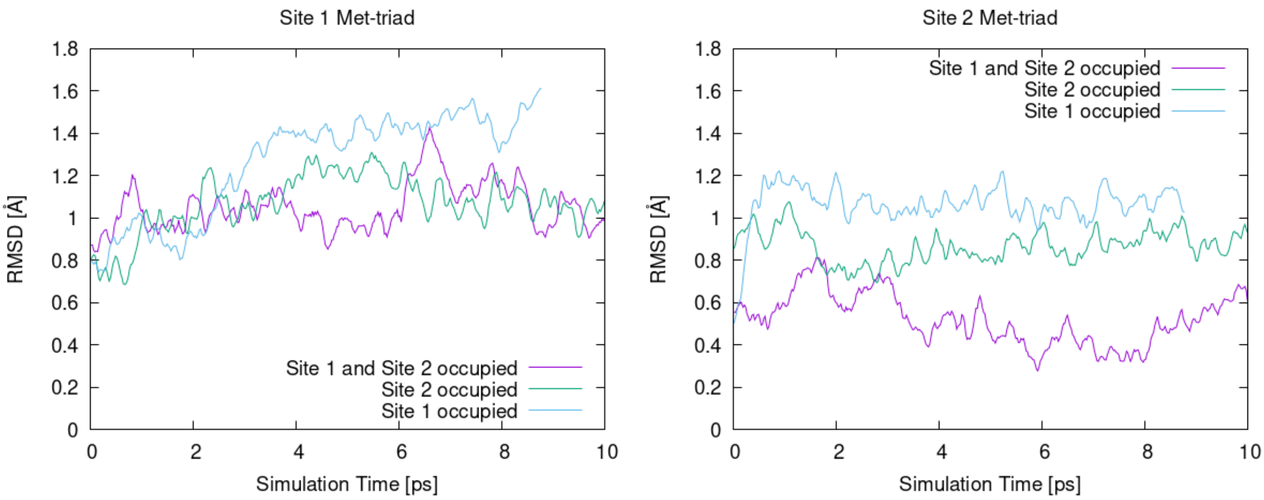

**Figure S4.** Root Mean Square Deviation (Å) RMSD vs simulation time (ps) of Site 1- (A) and Site 2- (B) Met-triads with respect to the initial conformation as obtained from QM/MM MD simulations. The CTR1 model with two Cu(I) ions, with Cu(I) bound to Site1 and with Cu(I) bound to Site 2 are shown in magenta, light blue and green lines, respectively.

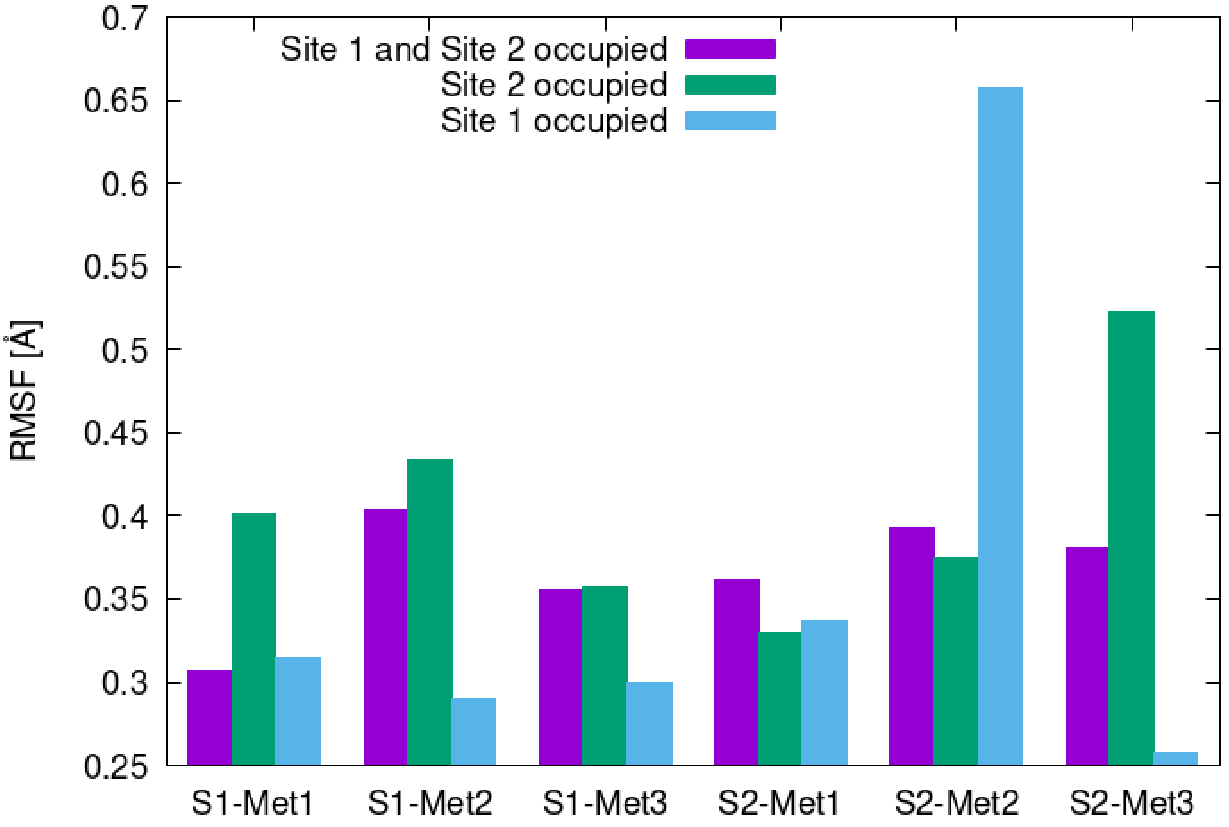

**Figure S5.** Root Mean Square Fluctuation (RMSF, Å) of the Met residues forming the Site 1 (S1-Met1-3) and Site 2 (S2-Met1-3) Met-triads. The CTR1 model with two Cu(I) ions, with Cu(I) bound to Site1 and with Cu(I) bound to Site 2 are shown in magenta, light blue and green lines, respectively.

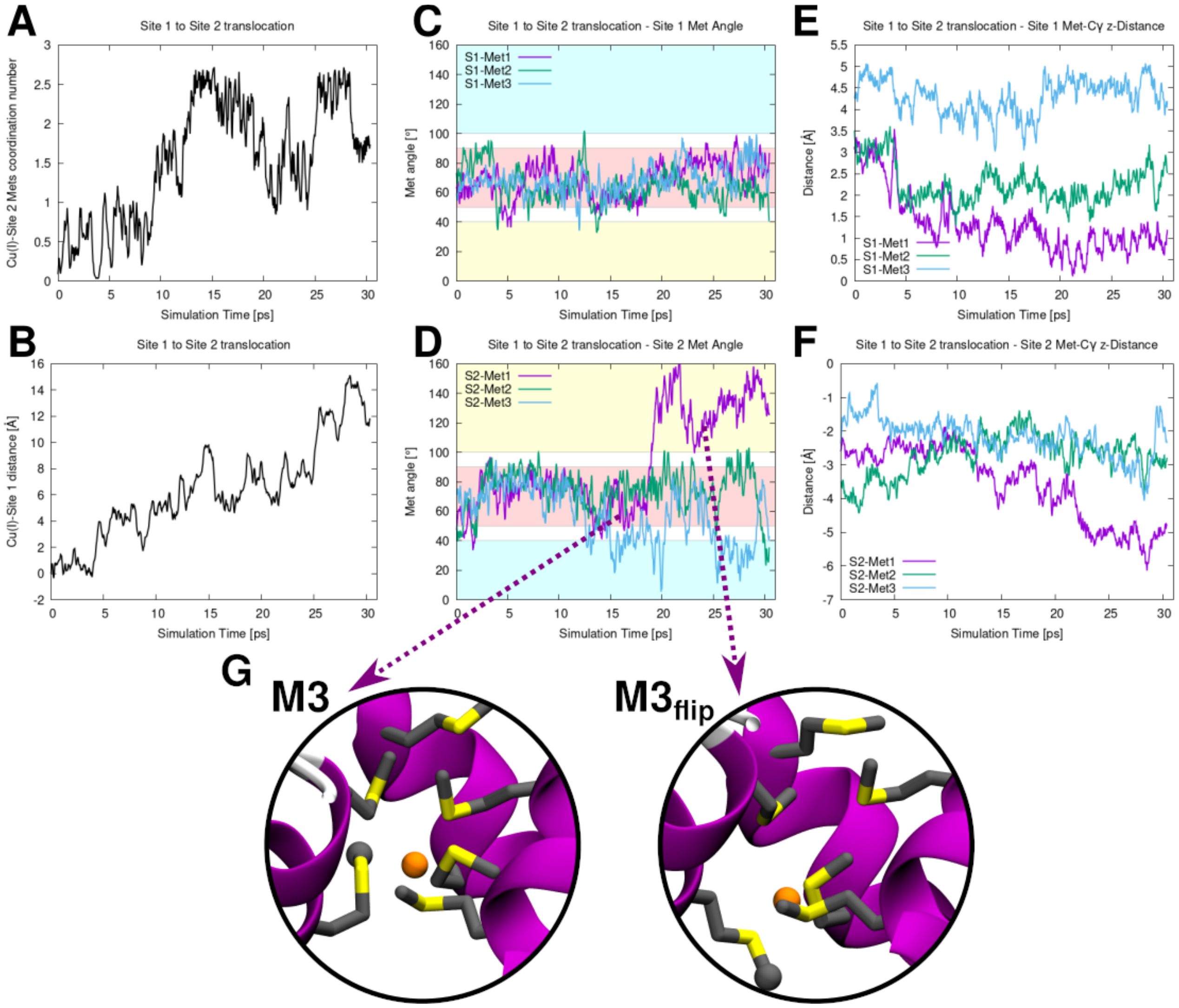

**Figure S6.** Cu(I) translocation from Site 1 to Site 2. Evolution of the Collective variables (CVs) used in the Metadynamics simulations vs simulation time (ps): (A) Coordination number of the C(I) ion to the S atoms of the Site 2 Met triad; (B) Distance of the Cu(I) to the S-Met atoms of the Site 1-Met triad; Conformations of the Site 1 (C) and Site 2 (D) Mets measured as the θ angle (°) between the Met Cγ-Cε vector and the vector of the selectivity filtered defined by geometric centers of Site 1 and Site 2 backbone atoms (see Figure 1C of the main text). S1-Met1-3 and S2-Met1-3 refer to the methionines of the top (extracellular matrix-exposed Met-triad) and bottom (intracellular-exposed Met-triad), respectively. Areas of the plots corresponding to the inward, in-plane and outward conformations are highlighted in cyan, red and yellow, respectively. Z-projection of distance (Å) of the Site 1 (E) and Site 2 (F) Met@Cγ to the center of the selectivity filter defined as the center of Site 1 and Site 2 Met@Cα atoms. (G) Close-ups of the states corresponding to the conformational change of the Site 2-Met1.

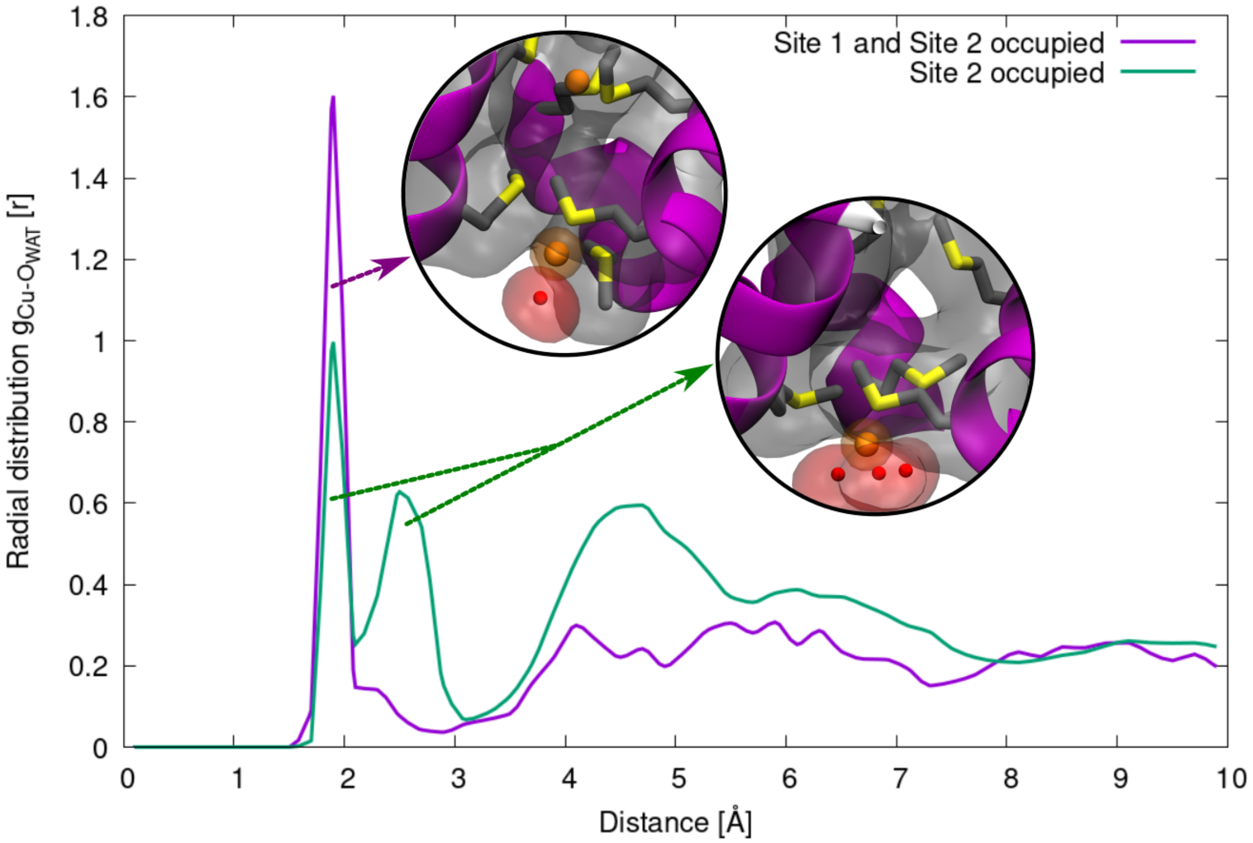

**Figure S7.** Radial distribution function (RDF) of the Cu(I) ions (bound to Site 2) – $O_{WAT}$ (oxygen atom of water molecules) distance in the model where both Site 1 and Site 2 bind a metal ion (purple line) and when only Site 2 binds a Cu(I) ion (green line). Inlaid pictures show close-up of the corresponding CTR1 conformations. CTR1 is shown as magenta new cartoons, the Met-triads are depicted as licorice and Cu(I) and water oxygen atoms as orange and red spheres, respectively. Hydrogens are omitted for clarity. CTR1 is highlighted as gray surface while water molecules and Cu(I) as red and orange surface, respectively.

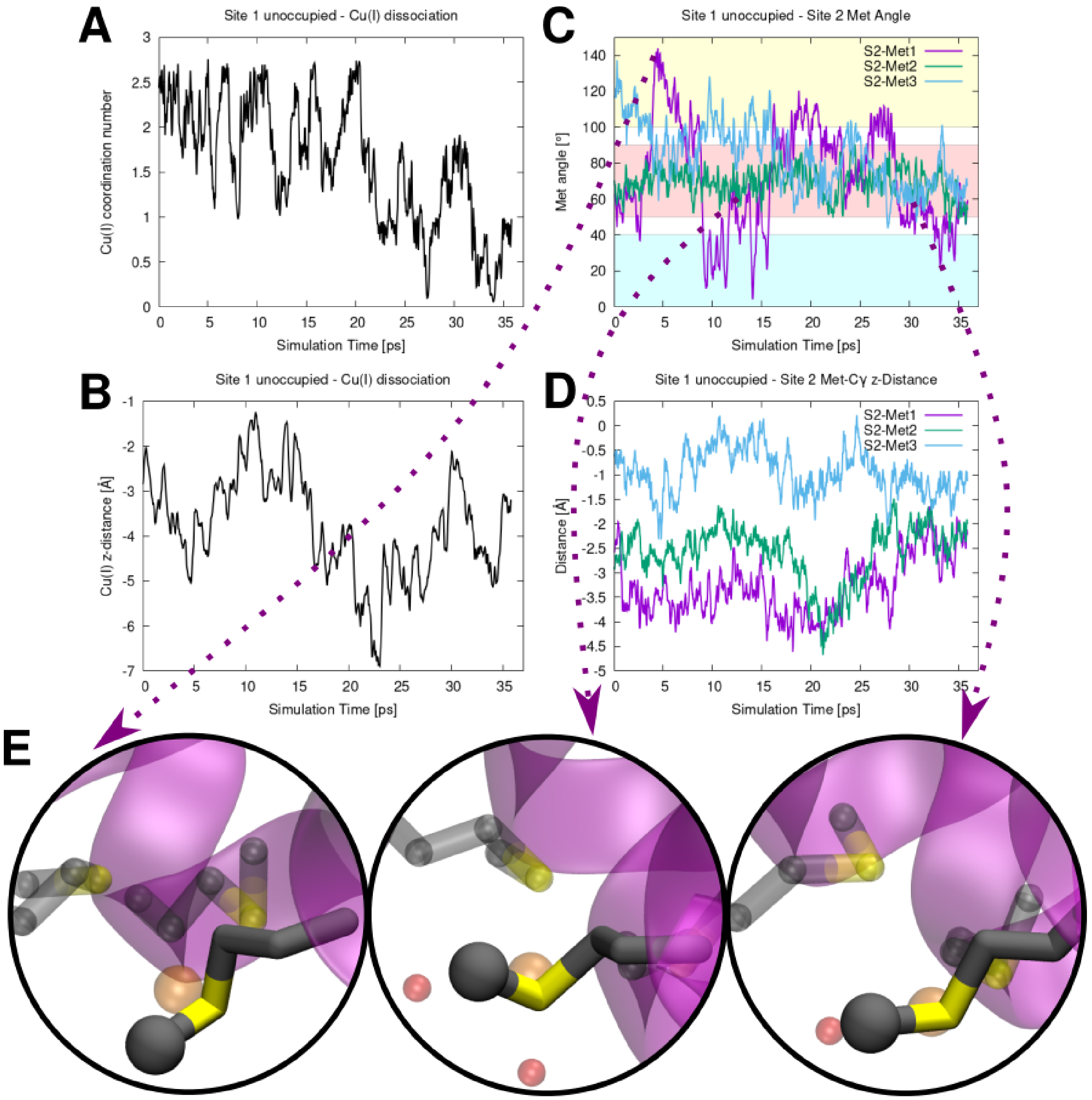

**Figure S8.** Cu(I) dissociation from Site 2 (S2) in the absence of Cu(I) bound to Site 1. Evolution of the Collective variables (CVs) used in the metadynamics simulation: (A) Coordination number of the Cu(I) ion to the S atoms of the S2-Met triad; (B) projection along the selectivity filter axis (z-axis) of the distance (Å) between the Cu(I) to the center of the filtration layer (defined as the center of of the S2-Met triad Cα atoms). (C) Conformation of the Site 2 Mets measured as the θ angle (°) between the Met Cγ-Cε vector and the vector of the selectivity filtered defined by geometric centers of Site 1 and Site 2 backbone atoms. S2-Met1-3 refer to the methionines of the bottom (intracellular-exposed Met-triad). Areas of the plot, corresponding to the inward, in-plane and outward conformations are highlighted in cyan, red and yellow, respectively. (D) Z-projection of distance (Å) of the Site 2 Met@Cγ to the center of the selectivity filter defined as the center of Site 1 and Site 2 Met@Cα atoms. (E) Close-ups of the states corresponding to the conformational change of the Site 2-Met1.

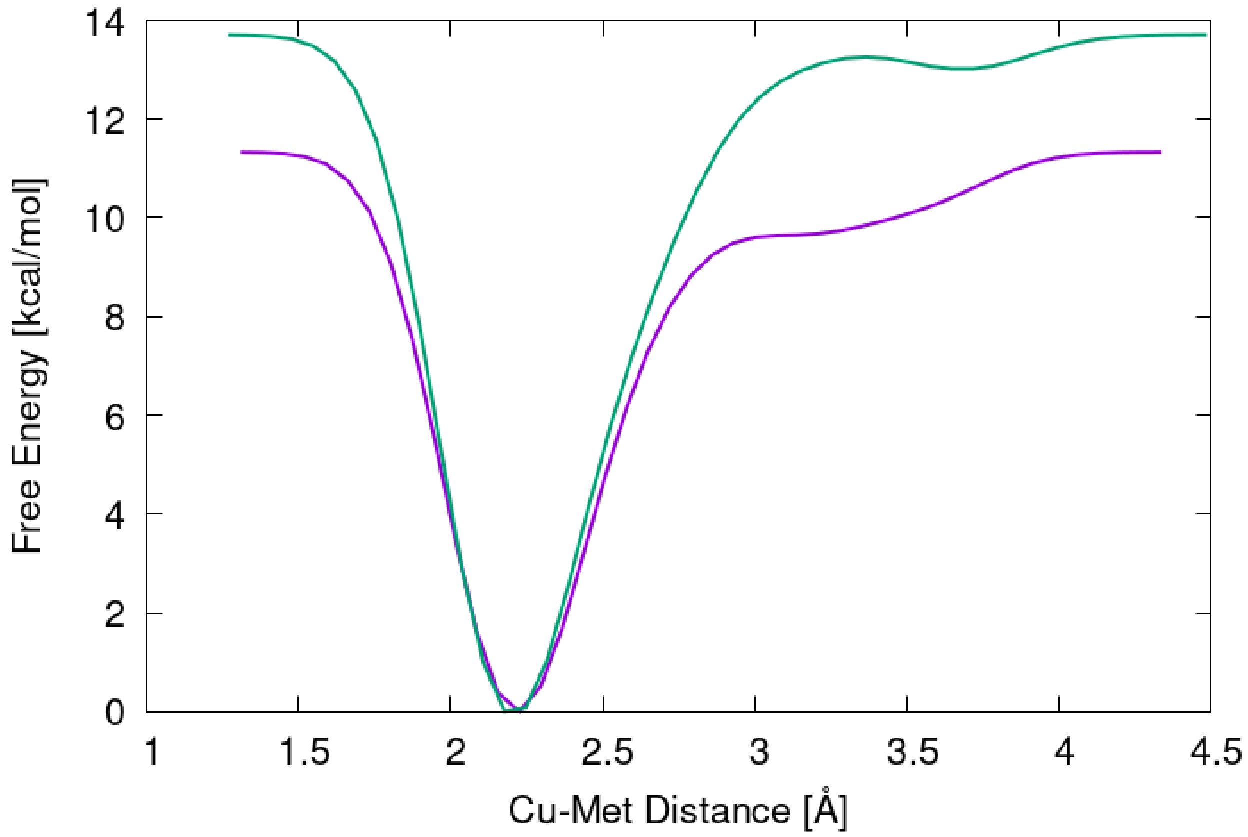

**Figure S9.** Free energy profile (kcal/mol) of the Cu(I) dissociation from Site 2 in absence of Cu(I) in Site 1 as a function of Cu(I) distance (Å) to the last coordinating Met@S atoms from the two repeats of the simulation (green and purple line).

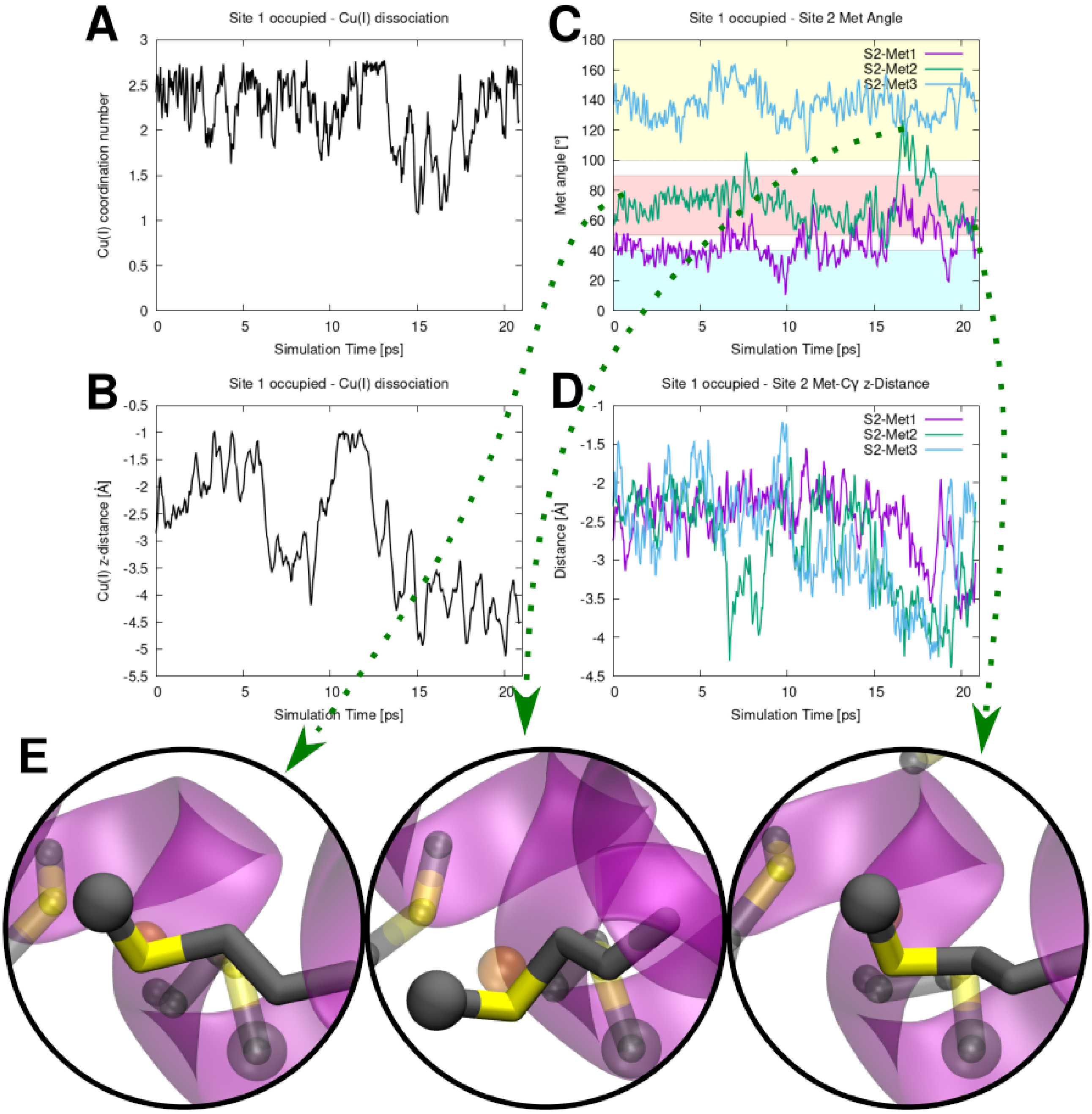

**Figure S10.** Cu(I) dissociation from Site 2 with a Cu(I) ion bound to Site 1. Evolution of the Collective variables (CVs) used in the metadynamics simulation: (A) Coordination number of the Cu(I) ions to the S atoms of the Site 2-Met triad; (B) projection along the selectivity filter axis (z-axis) of the distance (Å) between the Cu(I) to the center of the filtration layer (defined as the center of the Site 2-Met triad Cα atoms). (C) Conformation of the Site 2 Mets measured as the θ angle (°) between the Met Cγ-Cε vector and the vector of the selectivity filtered defined by geometric centers of Site 1 and Site 2 backbone atoms. S2-Met1-3 refer to the methionines of the bottom (intracellular-exposed Met-triad). Areas of the plot corresponding to the inward, in-plane and outward conformations are highlighted in cyan, red and yellow, respectively. (D) Z-projection of distance (Å) of the Site 2 Met@Cγ to the center of the selectivity filter defined as the center of Site 1 and Site 2 Met@Cα atoms. (E) Close-ups of the states corresponding to the conformational change of Site 2-Met2.

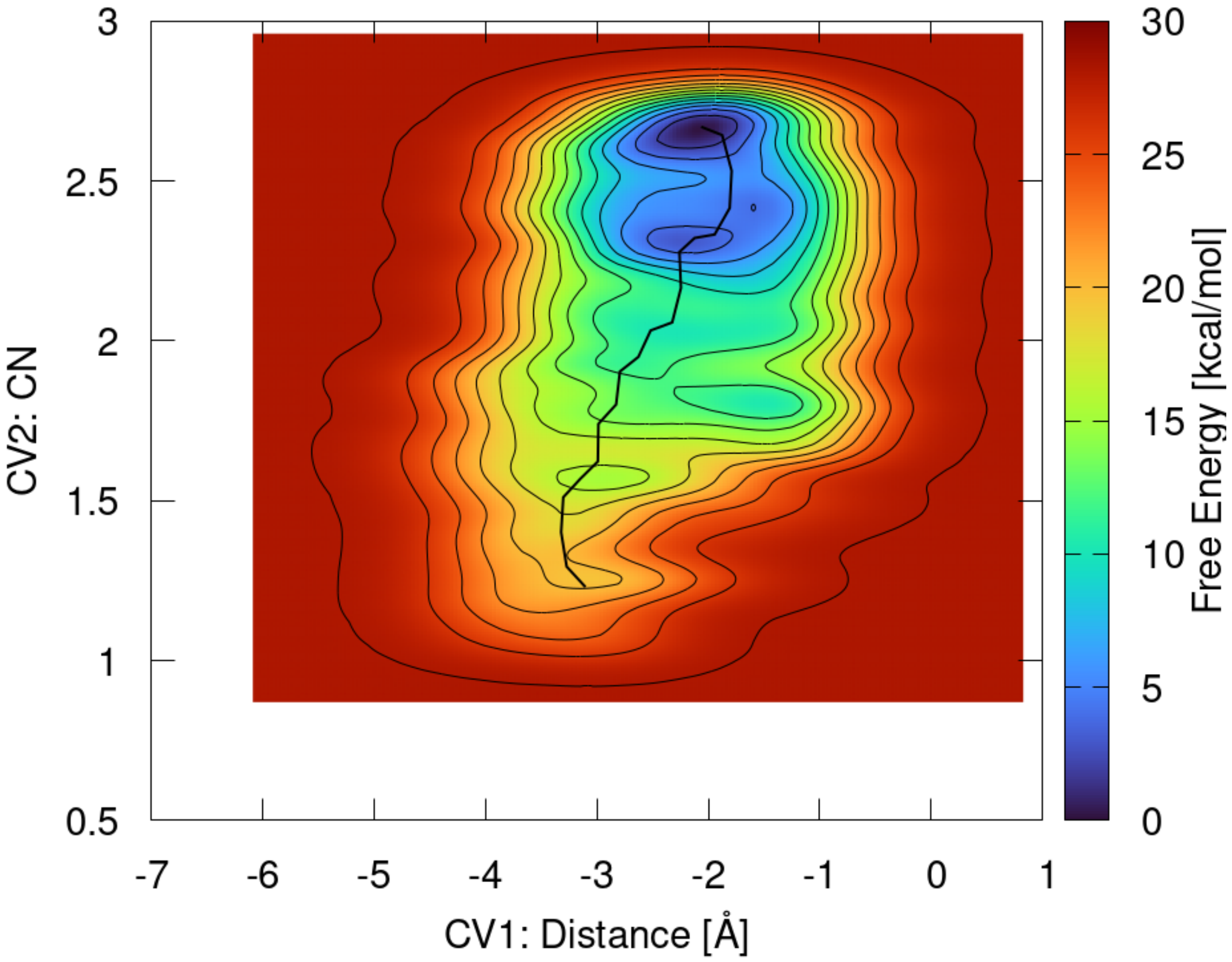

**Figure S11.** Free energy surface (FES, kcal/mol) of Cu(I) dissociation from Site 2 in the presence of Cu(I) ion bound to Site 1 and with the Site 2 Met-triads restrained to "in-plane" conformation. The FES is plotted as a function of the two Collective Variable (CVs) used in the metadynamics simulation: CV1 – projection along the z-axis of the distance between the Cu(I) ion and the center of Met-Cα atoms of the Site 2-Met triad; and CV2 – coordination number of Cu(I) with respect to the Site 2-Met triad S atoms. Minimum free energy path is plotted as a black line. The FES is shown from blue to red with isosurface lines drawn every 2.0 kcal/mol.

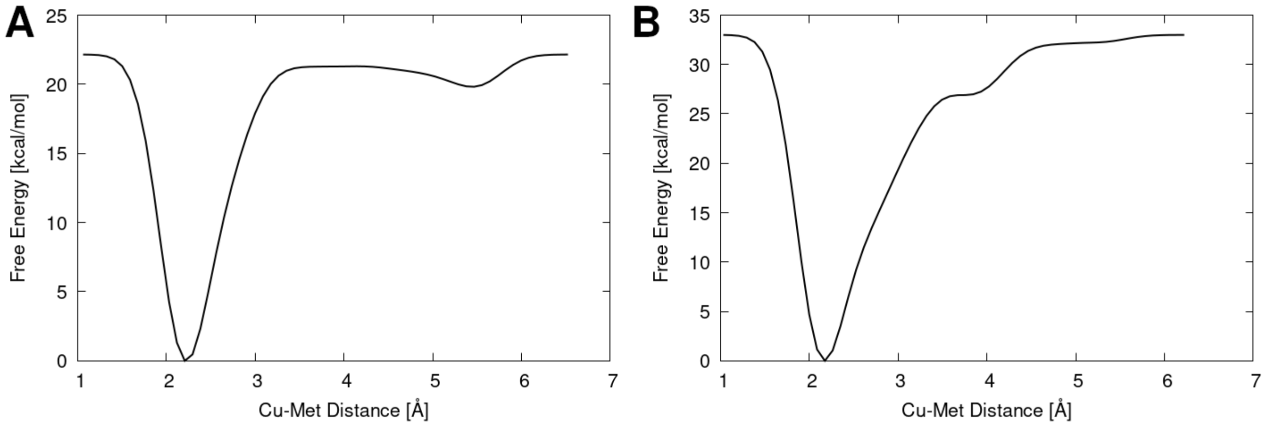

**Figure S12.** Free energy profiles (FEP, kcal/mol) of the final Cu(I) dissociation step from Site 2 in presence of Cu(I) in Site 1. The FEP is plotted as a function of the distance of the Cu(I) to the last coordinated Met@S atoms of Site 2. (A) All Mets restrained to the IP-conformation. (B) No restraints on the Mets conformation (i.e. one Met is in the O-conformation).

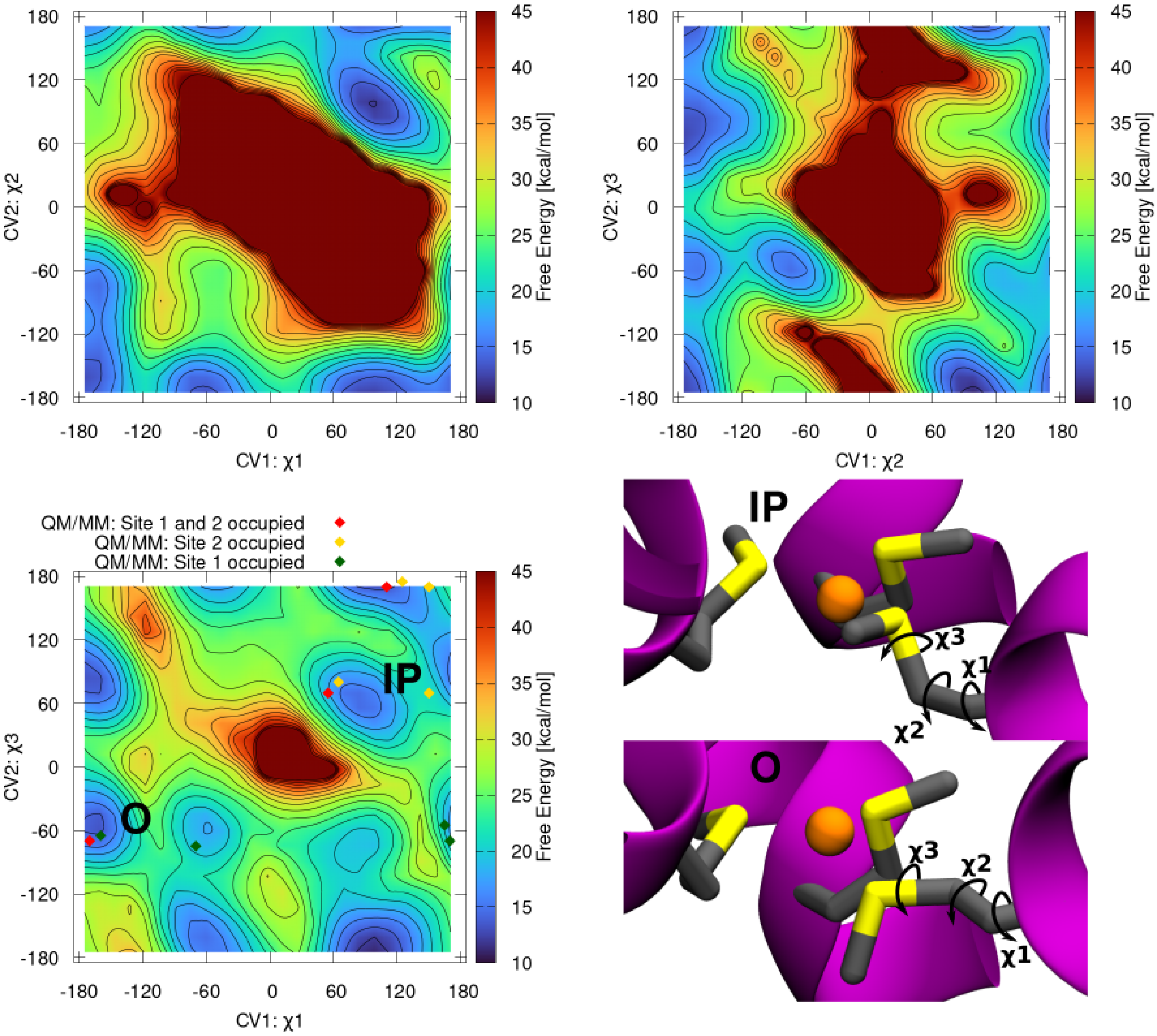

**Figure S13.** Free energy surfaces (FESs, kcal/mol) of a Site 2 Met conformational landscape as a function of the three torsion angles χ1, χ2 and χ3 as obtained from classical metadynamics simulation in the presence of a Cu(I) ion bound to both Site 1 and Site 2. In this simulation the other

S2-Mets are restrained into the IP-conformation. The FESs are shown from blue to red with isosurface lines drawn every 2.0 kcal/mol. The IP- and O-conformations can be described in the FES vs the $\chi1$, $\chi3$ surface, as the flipping involves mainly these two variables. Predominant conformations of the Site 2-Mets observed in QM/MM MD simulation for the CTR1 model in the presence of two Cu(I) ions and the models with single Cu(I) bound to either Site 1 and Site 2 are reported as red, green and yellow dots, respectively.
